# Petabase-scale sequence alignment catalyses viral discovery

**DOI:** 10.1101/2020.08.07.241729

**Authors:** Robert C. Edgar, Jeff Taylor, Victor Lin, Tomer Altman, Pierre Barbera, Dmitry Meleshko, Dan Lohr, Gherman Novakovsky, Benjamin Buchfink, Basem Al-Shayeb, Jillian F. Banfield, Marcos de la Peña, Anton Korobeynikov, Rayan Chikhi, Artem Babaian

**Author notes:** All authors contributed equally to this work.

## Abstract

Public databases contain a planetary collection of nucleic acid sequences, but their systematic exploration has been inhibited by a lack of efficient methods for searching this corpus, now exceeding multiple petabases and growing exponentially [1, 2]. We developed a cloud computing infrastructure, `Serratus`, to enable ultra-high throughput sequence alignment at the petabase scale. We searched 5.7 million biologically diverse samples (10.2 petabases) for the hallmark gene RNA dependent RNA polymerase, identifying well over 10^5^ novel RNA viruses and thereby expanding the number of known species by roughly an order of magnitude. We characterised novel viruses related to coronaviruses and to hepatitis *δ* virus, respectively and explored their environmental reservoirs. To catalyse a new era of viral discovery, we established a free and comprehensive database of these data and tools. Expanding the known sequence diversity of viruses can reveal the evolutionary origins of emerging pathogens and improve pathogen surveillance for the anticipation and mitigation of future pandemics.

## Introduction

Viral zoonotic disease has had a major impact on human health over the past century despite dramatic advances in medical science, notably the 1918 Spanish influenza, AIDS, SARS, Ebola, and COVID-19. There are an estimated 3×10^5^ mammalian virus species [3] from which infectious diseases in humans may arise [4], of which only a fraction is currently known. Global survey and monitoring of virus diversity is required for improved prediction and prevention of future epidemics; this effort is a focus of international consortia and hundreds of research laboratories worldwide [5, 6].

Virus discovery can be aided through re-analysis of the petabases of high-throughput sequencing data available in public databases such as the Sequence Read Archive (SRA) [1, 7–12], This data spans millions of ecologically diverse biological samples, many of which capture viral transcripts incidental to the goals of the original studies [13]. To catalyse global virus discovery, we developed the `Serratus` cloud computing infrastructure for ultra-high throughput sequence alignment.

We screened 5.7 million libraries comprising 10.2 petabases (10.2×10^15^ bases) of sequencing data, and report 883,502 RNA-dependent RNA polymerase (RdRP)-containing contigs which cluster at 90% amino-acid identity into 140,208 species-like operational taxonomic units (sOTUs), of which 132,260 were novel, i.e. >10% diverged from a previously known RdRP. We assembled 11,120 runs identified in this screen, recovering coronavirus and corona-like virus contigs including sequences from nine novel sOTUs. To demonstrate the broader utility of our approach, we also report 53 novel delta- and delta-like viruses related to the human pathogen hepatitis *δ* virus, and 252 representatives of a recently characterised family of huge bacteriophages.

Virus discovery is a fundamental step in preparing for the next pandemic. We lay the foundations for years of future research by enabling direct access to hundreds of thousands of virus sequences, captured from the collective efforts of over a decade of high-throughput sequencing studies. A free, interactive repository of `Serratus` data is available at https://serratus.io.

## Results

### Accessing the planetary virome with petabase-scale alignment

`Serratus` is a free, open-source cloud-computing infrastructure optimised to enable petabase-scale sequence alignment against a set of query sequences. Using `Serratus`, we aligned in excess of one million short-read sequencing datasets per day for under 1 US cent per dataset (Extended Figure 1). We used a widely available commercial computing service to deploy up to 22,250 virtual CPUs simultaneously (see Methods), leveraging SRA data mirrored onto cloud platforms as part of the NIH STRIDES initiative [14].

Our search space spans data deposited over 13 years from every continent and ocean, and all kingdoms of life (Figure 1). We applied `Serratus` in two of many possible configurations. First, to identify libraries containing known or closely related viruses we searched 3,837,755 (ca. May 2020) public RNA-seq, meta-genome, meta-transcriptome, and meta-virome datasets (termed sequencing runs [1]) against a nucleotide pangenome of all known coronavirus sequences plus all vertebrate RefSeq viruses (Extended Figure 3). We then aligned 5,686,715 runs (ca. January 2021) against known viral RdRP amino acid sequences, completing this search within 11 days (Figure 1a and Methods).

**Figure 1:**
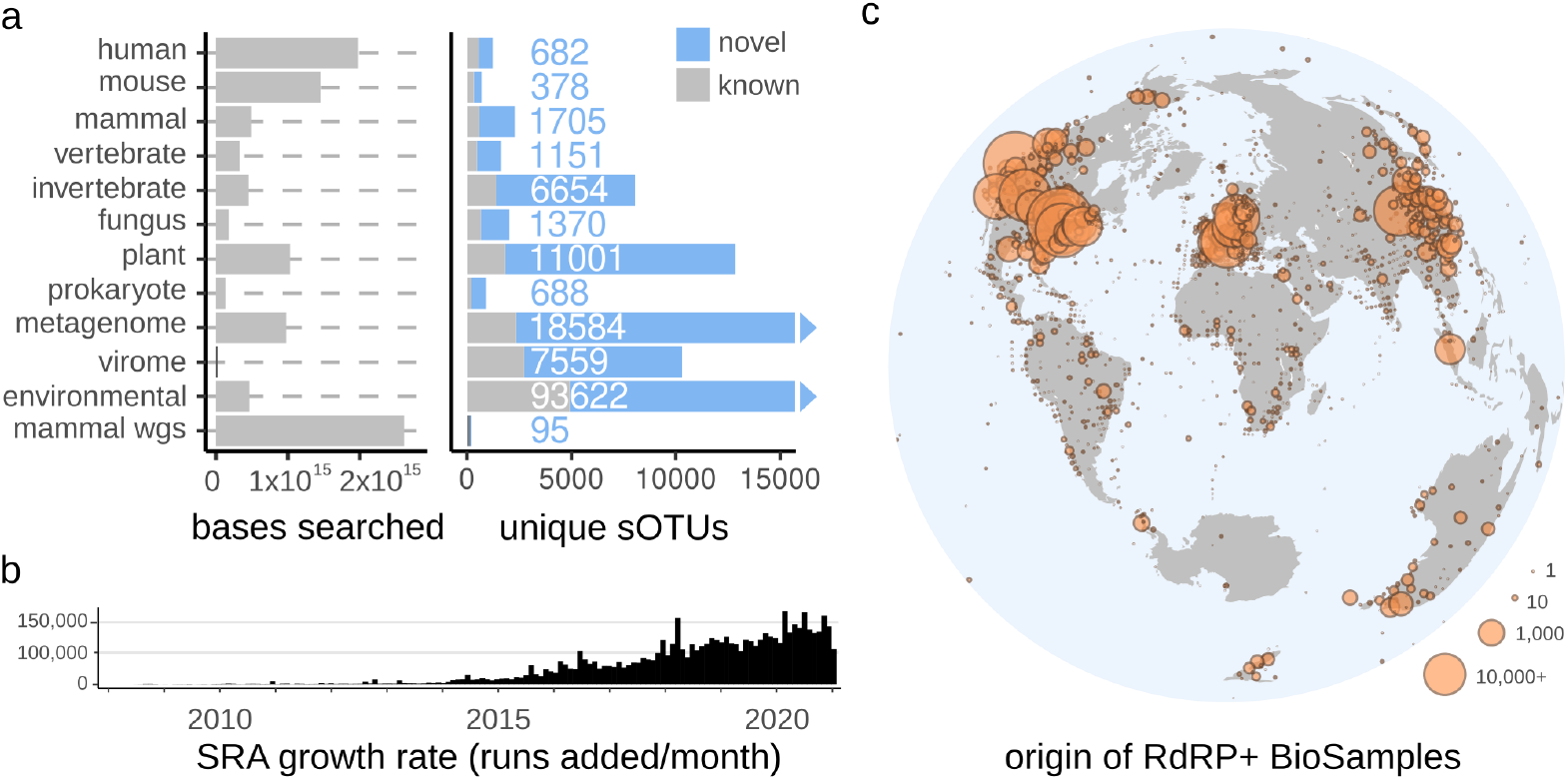
Searching the planetary virome. **a** Total bases searched from the 5,686,715 SRA sequencing runs analysed in the viral RdRP search grouped by sample taxonomy, where available (see Extended Figures 1 and 3, and Extended Table 1). 8,871/15,016 (59%) of known RdRP species-like operational taxonomic units (sOTUs) were observed in the SRA, and 131,957 unique and novel RdRP sOTUs were identified (see Extended Figure 2). sOTUs identified in multiple taxonomic groups are counted in each group separately, numbers shown indicate the number of novel sOTUs in each group. **b** Release dates of the runs included in the analysis reflecting the growth rate of available data. **c** Sample locations for 635,656 RdRP-containing contigs (27.8% of samples lacked geographic metadata). The high density of RdRP seen in North America, Western Europe and Eastern Asia reflects the substantial acquisition bias for samples originating from these regions. Interactive map is available at https://serratus.io/geo.

Previous approaches for identifying sequences across the entire SRA rely on pre-computed indexes [15–17] requiring exact substring or hash-based matches which limit sensitivity to diverged sequences (Extended Figure 1f). Pre-assembled reads (e.g. NCBI Transcriptome Shotgun Assembly database [2]) enable efficient alignment-based searches, [8], but are currently available only for a small fraction of the SRA. `Serratus` aligns a query of up to hundreds of Mb against unassembled libraries, achieving much greater sensitivity to diverged viruses compared to substring (k-mer) indexes while using far less computational resources than assembly (Figure 1g, and Methods).

### A sketch of viral RNA dependent RNA polymerase

Viral RdRP is a hallmark gene of RNA viruses which lack a DNA stage of replication [18]. We identified RdRP by a well-conserved amino acid sub-sequence we call the “palmprint”. Palmprints are delineated by three essential motifs which together form the catalytic core in the RdRP structure (Figure 2 and [19]). We constructed species-like operational taxonomic units (sOTUs) by clustering palmprints at a threshold of 90% amino-acid identity, chosen to approximate taxonomic species [19].

**Figure 2:**
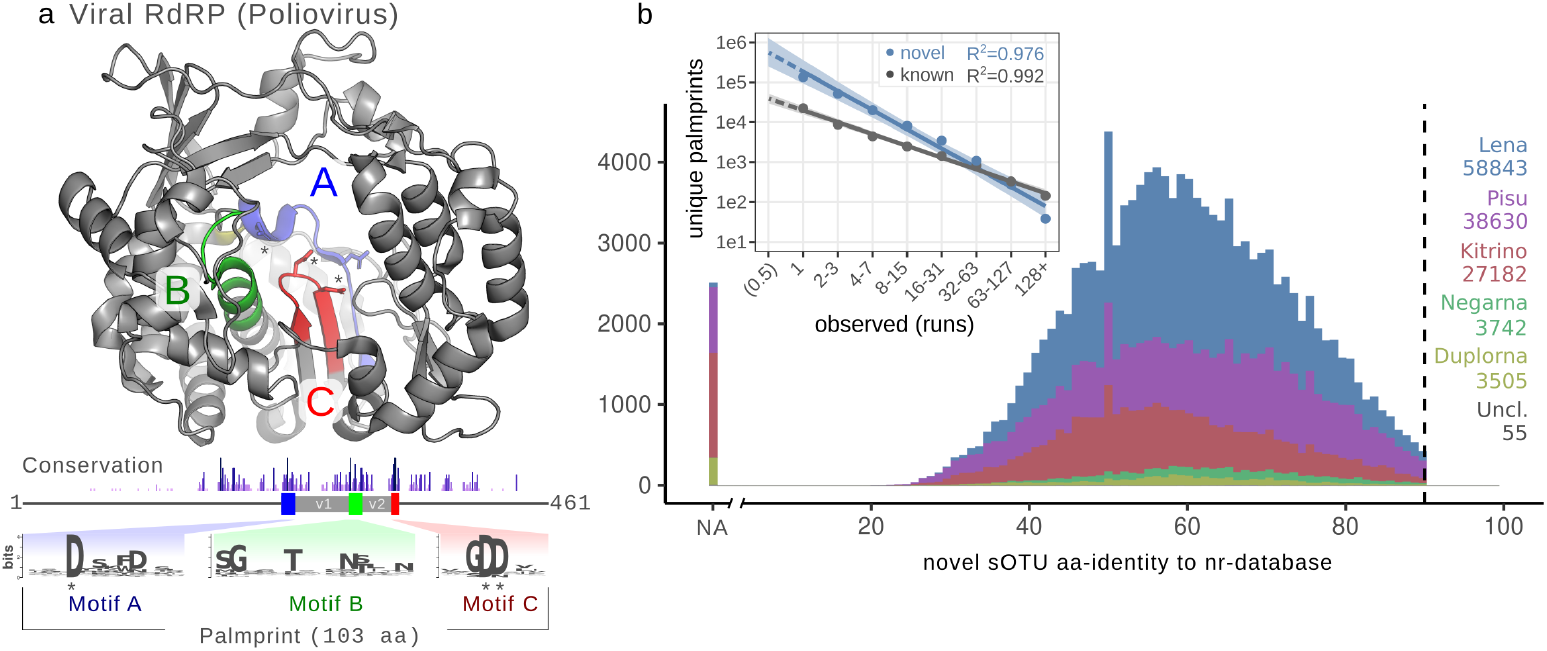
RNA dependent RNA Polymerase in the Sequence Read Archive. **a** The RdRP palmprint is the protein sequence spanning three well-conserved sequence motifs (A, B, and C), including intervening variable regions, exemplified within full-length poliovirus RdRP structure with essential aspartic acid residues(*) (pdb: 1RA6 [20]). Conservation was calculated from RdRP alignment in [21], trimmed to the poliovirus sequence; motif sequence logos are shown below. **b** Per-phylum histogram of amino acid identity of novel species-like operational taxonomic units (sOTUs) aligned to the NCBI non-redundant protein database. **inlay** Preston plot of palmprint abundances indicates that singleton palmprints (i.e., observed in exactly one run) occur within 95% confidence intervals of the value predicted by extrapolation from high-abundance palmprints (linear regression applied to log-transformed data), and this distribution is consistent through time (Extended Figure 2).

3,376,880 (59.38%) sequencing runs contained ≥1 reads mapping to the RdRP query (E-value ≤1e-4). We assembled aligned reads from each library (and their mate-pairs when available), yielding 4,261,616 contigs. 881,167 (20.7%) contained a high-confidence `palmprint` identified by Palmscan (false discovery rate = 0.001, further discussed in a companion manuscript [19]), representing 260,808 unique palmprints. Applying `Palmscan` to reference sequence databases[2, 21, 22] we obtained 45,824 unique palmprints, which clustered into 15,016 known sOTUs. If a newly acquired palmprint aligned to a known palmprint at ≥90% identity, it was assigned membership to that reference sOTU, otherwise it was designated novel. We clustered novel palmprints at 90% identity, obtaining 131,957 novel sOTUs, representing an increase of known RNA viruses by a factor ~9.8. Clustering novel palmprints at genus-like 75% and family-like 40% thresholds yielded 78,485 and 3,599 novel OTUs, representing increases of 8.0x and 1.9x, respectively (Figure 2b).

We extracted host, geospatial, and temporal metadata for each biological sample when available (Figure 1c), noting that the majority (88%) of novel RdRP sOTUs were observed from metagenomic or environmental runs, where accurate host inference is challenging. Mapping observations of virus marker genes across time and space suggests ecological niches for these viruses, while improved characterisation of sequence diversity can improve PCR primer design for *in situ* virus identification.

We estimate that ~1% of sOTUs are endogenous virus elements (EVEs), i.e. viral RdRPs which have serendipitously reverse-transcribed into a host germline. We did not attempt to systematically distinguish EVEs from virus RdRPs, noting that EVEs with intact catalytic motifs are likely to be recent insertions which can serve as a representative sequence for related exogenous viruses.

Most (60.5%) recovered palmprints were found in exactly one run (singletons), and are observed within the expected frequency range predicted by extrapolating from more abundant sequences (Figure 2b). The abundance distribution of distinct palmprints is consistent with log-log-linear for each year from 2015 to 2020 (Extended Figure 2e), and over time, singletons are confirmed by subsequent runs at an approximately constant rate (Extended Figure 2g). The majority of novel viruses will be singletons until the diversity represented by the search query and the fraction of the planetary virome sampled in the SRA both approach saturation. Extrapolating one year forward, when the SRA is expected to double in size, we project 430,000 (95% CI [330K, 561K]) additional unique palmprints will be identified by running `Serratus` with its current query (Figure 2b).

The total number of virus species is estimated to be 10^8^ to 10^12^ [23], thus our expansion captured at most 0.1% of the global virome. However, if exponential data growth continues, we are at the cusp of identifying a significant fraction of Earth’s total genetic diversity with tools such as `Serratus`.

### Expanding the scope of known Coronaviridae

The SARS-CoV-2 pandemic has significantly impacted human society. We further exemplify the potential of `Serratus` for virus discovery with *Coronaviridae* (CoV), including a recently proposed sub-family [24] which contains a CoV-like virus, Microhyla alphaletovirus 1 (MLeV), in the frog *Microhyla fissipes*, and Pacific salmon nidovirus (PsNV) described in the endangered *Oncorhynchus tshawytscha* [25].

First, we identified 52,772 runs containing ≥ 10 CoV-aligned reads or ≥ 2 CoV k-mers (32-mer, [17]). These runs were *de novo* assembled with a new version of synteny-informed `SPAdes` called `coronaSPAdes` (discussed in a companion manuscript [26]). This yielded 11,120 identifiable CoV contigs which we annotated for a comprehensive assemblage of *Coronaviridae* in the SRA (see Methods for discussion). With this training data we defined a scoring function to predict subsequent success of assembly (Extended Figure 3c).

CoV and neighbouring palmprints comprise 70 sOTUs, 44 of which are described in public databases. 17 CoV sOTUs contained partial RdRP (inclusive of full palmprint) from an amplicon-based virus discovery study not yet publicly deposited [27]. The remaining 9 sOTUs are novel viruses, with protein domains consistent with a CoV or CoV-like genome organisation (Extended Figure 4).

We operationally designate MLeV, PsNV and the nine novel viruses broadly as group *E*, noting that all were found in samples from non-mammalian aquatic vertebrates (Figure 3). Notably, *Ambystoma mexicanum* (axolotl) nidovirus (AmexNV) was assembled in 18 runs, 11 of which yielded common ~19 kb contigs. Easing the criteria of requiring an RdRP match in a contig, 28/44 (63.6%) of the runs from the associated studies were AmexNV positive [28–30]. Consistent assembly breakpoints in AmexNV, PsNV and similar viruses suggests that the viral genomes of this clade of CoV-like viruses are organised in at least two segments, one containing ORF1ab with RdRP, and a shorter segment containing a lamin-associated domain protein, spike and N’ accessory genes (Figure 3). An assembly gap with common breakpoints is present in the published PsNV genome [25]. Together these seven monophyletic species likely represent a distinct clade of segmented CoV-like nidoviruses, although molecular validation of this hypothesis is warranted.

**Figure 3:**
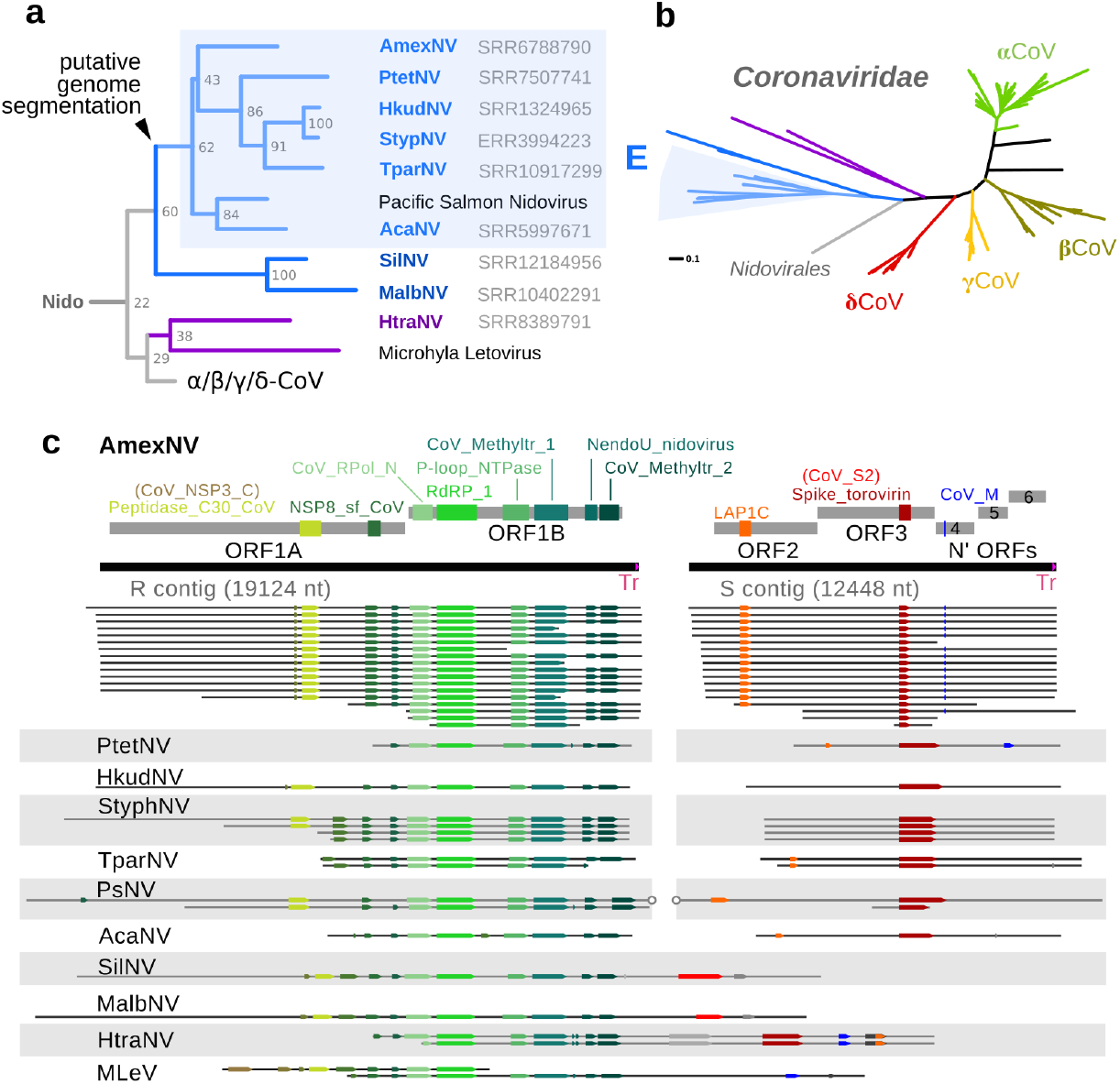
Expanding Coronaviridae. **a** Phylogram for group *E* sequences. Six viruses were similar to PsNV in *Ambystoma mexicanum* (axolotl; AmexNV), *Puntigrus tetrazona* (tiger barb; PtetNV), *Hippocampus kuda* (seahorse; HkudNV), *Syngnathus typhle* (broad-nosed pipefish; StypNV), *Takifugu pardalis* (fugu fish; TparNV), and the *Acanthemblemaria sp.* (blenny; AcaNV). More distant members identified were in *Hypomesus transpacificus* (the endangered delta smelt; HtraNV), *Silurus sp.* (catfish) SilNV, and *Monopterus albus* (asian swamp eel) MalbNV. **b** Unrooted phylogram for *Coronaviridae* annotated with genera (Greek letters) and group *E* CoV-like nidoviruses (see also Extended Figure 4). Maximum likelihood tree generated by clustering the RdRp amino acid sequences at 97% identity to show ~sub-species variability. **c** Genome structure of AmexNV and the contigs recovered from group *E* CoV-like viruses annotated with hidden-Markov model matches. AmexNV contigs contain an identical 129 nt trailing sequence (Tr). All the putatively segmented CoV-like are monophyletic with PsNV. A gap in the PsNV reference sequence [25] is shown with circles, overlapping the common contig ends seen in these viruses.

In addition to identifying genetic diversity within CoV, we cross-referenced CoV+ library meta-data to identify possible zoonoses and vectors of transmission. Discordant libraries, one in which a CoV is identified and the viral expected host [31] does not match the sequencing library source taxa, were rare, accounting for only 0.92% of cases (Extended Table 1e).

In data from a 2010 virome sequencing study of children with febrile illness [32], we identified sequencing runs from two children, one febrile (SRR057949) and one afebrile (SRR057961) with reads mapping to the *β*-CoV, Murine Hepatitis Virus (MHV). We assembled a complete 31.3 kb MHV genome from each replicate taken from the febrile child and a partial genome from the afebrile child. MHV can infect human cells *in vitro* [33], but clinical infection may be rare and missed by targeted diagnostic assays. Infectious agents are the leading cause of pyrexia of unknown origin (PUO) in children and immunocompromised adults [34]. This highlights the need for rapid and unbiased metagenomic sequence diagnostics, technically akin to `Serratus`. This would help resolve the etiology of a sub-set of PUO and serve as an early-warning surveillance system for zoonoses, enabling more efficacious public-health intervention.

An important limitation for these analyses is that the nucleic acid reads do not prove viral infection has occurred in the nominal host species. For example, we identified five libraries in which a porcine, avian or bat coronavirus was found in plant samples. The parsimonious explanation is that CoV was present in faeces/fertiliser originating from a mammalian or avian host applied to these plants. However, this exemplifies a merit of unbiased search in identifying transmission vectors and monitoring the geo-temporal boundaries of a virus.

### Rapid expansion into the viral unknowns

The global mortality from viral hepatitis exceeds that of HIV/AIDS, tuberculosis or malaria [35]. Hepatitis *δ* virus (HDV) has a small circular RNA genome (~1.7 knt) which folds into a rod-like shape and encodes three genes: a delta antigen protein, and two self-cleaving ribozymes (drbz) [36].

Prior to 2018, HDV was the sole known member of its genus; 13 drbz-containing members have since been characterised [37–42], and recently a second class of ribozyme (known as hammerhead or hhrbz) characteristic of plant viroids was identified in delta-like viruses we refer to as epsilon viruses [43]. By sequence search for the delta antigen protein and ribozymes, we identified 14 deltaviruses, 39 epsilonviruses and 311 enigmatic sequences with deltavirus-like synteny we term zeta viruses (Figure 4, Extended Figure 5). The evolutionary histories of these mammalian deltaviruses are explored further in a companion paper [41].

**Figure 4:**
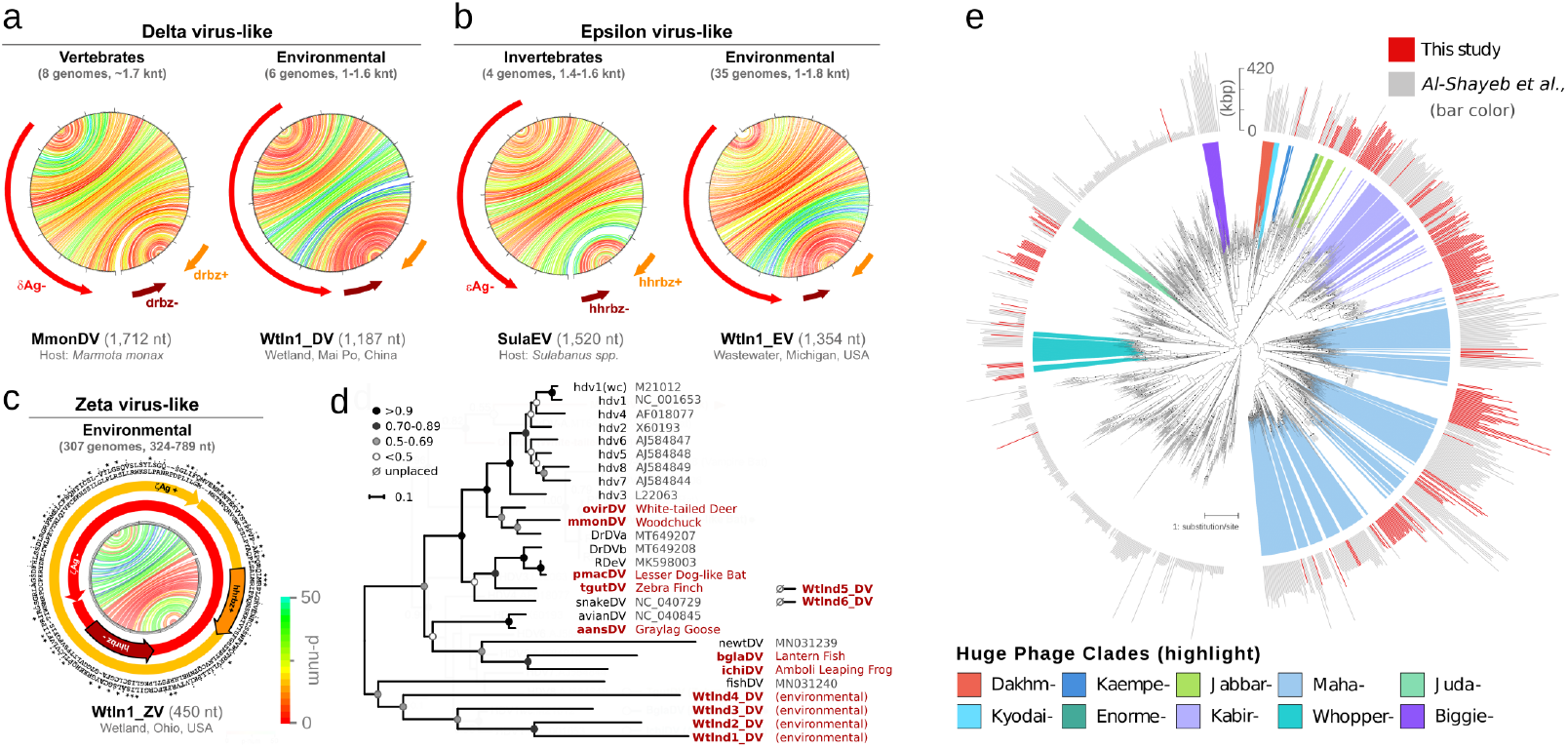
Expanding deltaviruses and hugephages. **a** Genome structure for the *Marmota monax* Delta virus (MmonDV) and a DV-like genome detected in an environmental dataset each containing a negative-sense delta-antigen (*δ*Ag) ORF; two delta ribozymes (dvrbz); and characteristic rod-like folding, where each line shows the predicted base-pairing within the RNA genome, coloured by base-pairing confidence score (p-num) [44]. **b** Similar genome structure for the *Sulabanus* spp. Epsilon virus-like (SulaEV) and an EV-like genome from an environmental dataset each containing a negative-sense epsilon-antigen (*ϵ*Ag) ORF; two hammerhead ribozymes (hhrbz); and rod-like folding. **c** Example of the compact genome structure of a Zeta virus-like from an environmental dataset containing two predicted zeta-antigen (*ζ*ZAg+/−, protein alignment is shown in the outer circles) ORFs without stop codons; two hhrbz overlapping with the ORFs; and rod-like folding. Further novel genomes are shown in Extended Figures 5 and 6. **d** Maximum-likelihood phylogenetic tree of DVs derived from a delta-antigen protein alignment with bootstrap values. Two divergent environmental DV could not yet be placed. **e** Tree showing huge phage clade expansion. Black dots indicate branches with bootstrap values >90. Outer ring indicates genome or genome fragment length: gray are sequences from [45] and reference sequences, shadings indicate previously defined clades of phages with very large genomes (200-735 kbp). The Kabirphages (light purple) are shown in expanded view in Extended Figure 7.

The zeta virus circular genomes are highly compressed, ranging from 324-789 nt and predicted to fold into rodlike structures. They contain a hhrbz in each orientation and encode two ORFs, one sense and one anti-sense. Both ORFs generally lack stop codons and encompass the entire genome, potentially producing an endless tandem-repeat of antigen. The atypical coiled-coil domain of the HDV antigen [46] is conserved in the antigens of new delta and epsilon viruses, whereas epsilon and zeta genomes show analogous hhrbzs (Extended Figure 6), supporting that these sequences may share common ancestry. These abundant elements may help to solve a long-standing question about the origins of circular RNA subviral agents in higher eukaryotes (Extended Figure 6), historically regarded as molecular fossils of a prebiotic RNA world [47]

To evaluate the feasibility of applying `Serratus` in the context of microbiome research, we sought to locate bacteriophages related to recently reported huge phages [45], searching for terminase amino acid sequences. Targeted assembly of 287 high-scoring runs returned 252 terminase-containing contigs ≥140 kbp. Phylogenetics of these sequences resolved new groups of phages with large genomes (Figure 4e). While most phages were from a single animal genus, we identified closely related phages crossing animal orders, including related phages in a human from Bangladesh (ERR866585) and groups of cats (PRJEB9357) and dogs (PRJEB34360) from England, sampled 5 years apart. Similarly, we recovered two 554 kbp Lak megaphage genomes (among the largest animal microbiome phages reported to date) that are extremely closely related to sequences previously reported from pigs, baboons and humans (Extrended Figure 7), [48]. These two genomes were circularised and manually curated to completion. The large carrying capacity of such phages and broad distribution underlines their potential for extensive lateral gene transfer amongst animal microbiomes and modification of host bacterial function. The newly-recovered sequences substantially expand and augment the inventory of phages with genomes whose length range overlaps with those of bacteria.

## Discussion

Since the completion of the the human genome, growth of DNA sequencing databases has outpaced Moore’s Law [49]. `Serratus` provides rapid and focused access to genomic sequences captured over more than a decade by the global research community which would otherwise be inaccessible in practice. This work and further extensions of petabase scale genomics [16, 17, 50] are shaping a new era in computational biology, enabling expansive gene discovery, pathogen surveillance, and pangemomic evolutionary analyses.

Optimal translation of such massive datasets into meaningful biomedical advances requires free and open collaboration amongst scientists [51, 52]. The current pandemic underscores the need for prompt, unrestricted and transparent data sharing. With these goals in mind, we deposited 7.3 terabytes of virus alignments and assemblies into an open-access database which can be explored via a graphical web interface at https://serratus.io or programatically through the `tantalus R` package and its `PostgreSQL` interface.

The “metagenomics revolution“ of virus discovery is accelerating [22, 53]. Innovative fields such as high-throughput viromics [54] can leverage vast collections of virus sequences to inform policies that predict and mitigate emerging pandemics [55, 56]. Combining ecoinformatics with virus, host, and geotemporal metadata offers a proof of concept for a global pathogen surveillance network, arising as a byproduct of centralised and open data sharing.

Human population growth and encroachment on animal habitats is bringing more species into proximity, leading to increased rate of zoonosis [4] and accelerating the Anthropocene mass extinction [57, 58]. `Serratus` enhances our capability to chronicle the full genetic diversity of our planet’s diminishing biosphere. The need to invest in collection and curation of biologically diverse samples with emphasis on geographically under-represented regions has never been more pressing. If not for the conservation of endangered species, then to better conserve our own.

## 1 Materials and Methods

### 1.1 Serratus alignment architecture

`Serratus` (https://github.com/ababaian/serratus) is an open-source cloud-infrastructure designed for ultra-high throughput sequence alignment against a query sequence or pangenome (Extended Figure 1). `Serratus` compute costs are dependent on search parameters (expanded discussion available: https://github.com/ababaian/serratus/wiki/pangenome_design). The nucleotide vertebrate viral pangenome search (`bowtie2`, database size: 79.8 Mb) reached processing rates of 1.29 million SRA runs in 24-hours at a cost of $0.0062 US dollars per dataset (Extended Figure 1). The translated-nucleotide RdRP search (`DIAMOND`, database size: 7.1 Mb) reached processing rates exceeding 0.5 million SRA runs in 12-hours at a cost of $0.0042 per dataset. All 5,686,715 runs analysed in the RdRP search were completed within 11 days for a total cost of $23,980 or 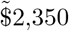 per petabase. For a detailed breakdown of `Serratus` project costs and recommendations for managing cloud-computing costs, see Serratus wiki: https://github.com/ababaian/serratus/wiki/budget.

#### 1.1.1 Computing cluster architecture

The processing of each sequencing library is split into three modules `dl` (download), `align`, and `merge`. The `dl` module acquires compressed data (.sra format) via prefetch, from the Amazon Web Services (AWS) Simple Storage Service (S3) mirror of the Sequence Read Archive (SRA), decompresses to FASTQ with `fastq-dump`, and splits the data into chunks of 1 million reads or read-pairs (`fq-blocks`) into a temporary S3 cache bucket. To mitigate excessive disk usage caused by a few large datasets, a total limit of 100 million reads per dataset was imposed. The `align` module reads individual fq-blocks and aligns to an indexed database of user-provided query sequences using either `bowtie2` (v2.3.4.1,`--very-sensitive-local`) [59] for nucleotide search, or `DIAMOND` (v2.0.6 - development version, `--mmap-target-index --target-indexed --masking 0 --mid-sensitive -s 1 -c1 -p1 -k1 -b 0.75`) [60] for translated-protein search. Finally, the `merge` module concatenates the aligned blocks into a single output file (`.bam` for nucleotide, or `.pro` for protein) and generates alignment statistics with a Python script (see `Summarizer` below).

#### 1.1.2 Computing resource allocation

Each component is launched from a separate AWS autoscaling group with its own launch template, allowing the user to tailor instance requirements per task. This enabled us to minimise the use of costly block storage during compute-bound tasks such as alignment. We used the following Spot instance types; `dl`: 250GB SSD block storage, 8vCPUs, 32GB RAM (r5.xlarge) ~1300 instances; `align`: 10GB SSD block storage, 8vCPUs, 8GB RAM (c5.xlarge) ~4,300 instances; `merge`: 150GB SSD block storage, 4vCPUs, 4GB RAM (c5.large) ~60 instances. Users should note that it may be necessary to submit a service ticket to access more than the default 20 EC2 instance limit.

AWS Elastic Compute Cloud (EC2) instances have higher network bandwidth (up to 1.25 GB/s) than block storage bandwidth (250 MB/s). To exploit this, we used S3 buckets as a data buffering and streaming system to transfer data between instances following methods developed in a previous cloud architecture (https://github.com/FredHutch/sra-pipeline). This, combined with splitting of FASTQ files into individual blocks, effectively eliminated file input/output (i/o) as a bottleneck, since the available i/o is multiplied per running instance (conceptually analogous to a RAID0 configuration or a `Hadoop` distributed filesystem [61].

Using S3 as a buffer also allowed us to decouple the input and output of each module. S3 storage is cheap enough that in the event of unexpected issues (e.g., exceeding EC2 quotas) we could resolve system problems in realtime and resume data processing. For example, shutting down the `align` modules to hotfix a genome indexing problem without having to re-run the `dl` modules, or if an alignment instance is killed by a Spot termination, only that block needs to be reprocessed instead of the entire sequencing run.

#### 1.1.3 Work queue and scheduling

The `Serratus` scheduler node controls the number of desired instances to be created for each component of the workflow, based on the available work queue. We implemented a pull-based work queue. Upon boot-up each instance launches a number of worker threads equal to the number of CPU available. Each worker independently manages itself via a boot script, and queries the scheduler for available tasks. Upon completion of the task, the worker updates the scheduler of the result: success, or fail, and queries for a new task. Under ideal conditions, this allows for a worst-case response rate in the hundreds of milliseconds, keeping cluster throughput high. Each task typically lasts several minutes depending on the pangenome.

The scheduler itself was implemented using Postgres (for persistence and concurrency) and Flask (to pool connections and translate REST queries into SQL). The Flask layer allowed us to scale the cluster past the number of simultaneous sessions manageable by a single Postgres instance. The work queue can also be managed manually by the user, to perform operations such as re-attempt downloading of an SRA accession upon a failure or to pause an operation while debugging. Up to 300,000 SRA jobs can be processed in the work queue per batch process.

The system is designed to be fully self-scaling. An ”autoscaling controller” was implemented which scales-in or scales-out the desired number of instances per task every five minutes based on the work queue. As a backstop, when all workers on an instance fail to receive work instructions from the scheduler, the instance self shuts-down. Finally a ”job cleaner” component checks the active jobs against currently running instances. If an instance has disappeared due to SPOT termination or manual shutdown, it resets the job allowing it to be processed up by the next available instance.

To monitor cluster performance in real-time, we used `Prometheus` and `node_exporter` to retrieve CPU, disk, memory, and networking statistics from each instance, `postgres_exporter` to expose performance information about the work queue, and Python exporter to export information from the Flask server. This allowed us to identify and diagnose performance problems within minutes to avoid costly overruns.

#### 1.1.4 Generating viral summary reports

We define a viral pangenome as the entire collection of reference sequences belonging to a taxonomic viral family, which may contain both full-length genomes and sequence fragments such as those obtained by RdRP amplicon sequencing.

We developed a `Summarizer` module written in Python to provide a compact, human- and machine-readable synopsis of the alignments generated for each SRA dataset. The method was implemented in `serratus_summarizer.py` for nucleotide alignment and `serratus_psummarizer.py` for amino acid alignments. Reports generated by the `Summarizer` are text files with three sections described in detail online (https://github.com/ababaian/serratus/wiki/.summary-Reports). In brief, each contains a header section with alignment meta-data and one-line summaries for each virus family pangenome, reference sequence and gene respectively, with gene summaries provided for protein alignments only.

For each summary line we include descriptive statistics gathered from the alignment data such as the number of aligned reads, estimated read depth, mean alignment identity, and coverage, i.e. the distribution of reads across each reference sequence or pangenome. Coverage is measured by dividing a reference sequence into 25 equal bins and depicted as an ASCII text string of 25 symbols, one per bin; for example `oaooomoUU:oWWUUWOWamWAAUW`. Each symbol represents ⌊*log_2_*(*n* + 1⌋ where *n* is the number of reads aligned to a bin in this order: 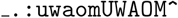. Thus, ‘_’ indicates no reads, ‘.’ exactly one read, ‘:’ two reads, ‘`u`’ 3-4 reads, ‘`w`’ 5-7 reads and so on; 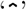 represents > 2^13^ = 8,192 reads in the bin. For a pangenome, alignments to its reference sequences are projected onto a corresponding set of 25 bins. For a complete genome, the projected pangenome bin number 1, 2,…, 25 is the same as the reference sequence bin number. For a fragment, a bin is projected onto the pangenome bin implied by the alignment of the fragment to a complete genome. For example, if the start of a fragment aligns half way into a complete genome, bin 1 of the fragment is projected to bin 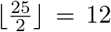 of the pangenome. The introduction of pangenome bins was motivated by the observation that `bowtie2` selects an alignment at random when there are two or more top-scoring alignments, which tends to distribute coverage over several reference sequences when a single viral genome is present in the reads. Coverage of a single reference genome may therefore be fragmented, and binning to a pangenome better assesses coverage over a putative viral genome in the reads while retaining pangenome sequence diversity for detection.

#### 1.1.5 Identification of viral families within a sequencing dataset

The `Summarizer` implements a binary classifier predicting the presence or absence of each virus family in the query. For a given family *F*, the classifier reports a score in the range [0,100] with the goal of assigning a high score to a dataset if it contains *F* and a low score if it does not. Setting a threshold on the score divides datasets into disjoint subsets representing predicted positive and negative detections of family *F*. The choice of threshold implies a trade-off between false positives and false negatives. Sorting by decreasing score ranks datasets in decreasing order of confidence that *F* is present in the reads.

Naively, a natural measure of the presence of a virus family is the number of alignments to its reference sequences. However, alignments may be induced by non-homologous sequence similarity, for example low-complexity sequence. The score for a family was therefore designed to reflect the overall coverage of a pangenome because coverage across all or most of a pangenome is more likely to reflect true homology, i.e. the presence of a related virus. Ideally, coverage would be measured individually for each base in the reference sequence, but this could add undesirable overhead in compute time and memory for a process which is executed in the Linux alignment pipe (FASTQ decompression → aligner → `Summarizer` → alignment file compression). Coverage was therefore measured by binning as described above, which can be implemented with minimal overhead.

A virus that is present in the reads with coverage too low to enable an assembly may have less practical value than an assembled genome. Also, genomes with lower identity to previously known sequences will tend to contain more novel biological information than genomes with high identity and will tend to have fewer alignments highly diverged segments. With these considerations in mind, the classifier was designed to give higher scores when coverage is high, read depth is high, and/or identity is low. This was accomplished as follows. Let *H* be the number of bins with at least 8 alignments to *F*, and *L* be the number of bins with from 1 to 7 alignments. Let *S* be the mean alignment percentage identity, and define the identity weight 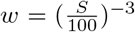, which is designed to give higher weight to lower identities, noting that *w* is close to one when identity is close to 100% and increases rapidly at lower identities. The classification score for family *F* is calculated as *Z_F_* = *max(w(4H* + L)), 100). By construction, *Z_F_* has a maximum of 100 when coverage is consistently high across a pangenome, and is also high when identity is low and coverage is moderate, which may reflect high read depth but many false negative alignments due to low identity. Thus, *Z_F_* is greater than zero when there is at least one alignment to *F* and assigns higher scores to SRA datasets which are more likely to support successful assembly of a virus belonging to *F*.

#### 1.1.6 Sensitivity to novel viruses as a function of identity

We aimed to assess the sensitivity of our pipeline as a function of sequence identity by asking what fraction of novel viruses is detected at increasingly low identities compared to the reference sequences used for the search. Several variables other than identity affect sensitivity, including read length, whether reads are mate-paired, sequencing error rate, coverage bias, and presence of other similar viruses which may cause some variants to be unreported in the contigs. Coverage bias can render a virus with high average read depth undetectable, in particular if the query is RdRP-only and the RdRP gene has low coverage or is absent from the reads. Successful detection might be defined in different ways, depending on the goals of the search; e.g. a single local alignment of a reference to a read (maximising sensitivity, but not always useful in practice); a micro-assembled palmprint; a full assembly contig that contains a complete palmprint or otherwise classifiable fragment of a maker gene; or an assembly of a complete genome. We assessed alignment sensitivity of `bowtie2 --very-sensitive-local` and Serratus-optimised `DIAMOND` as a function of identity by simulating typical examples in representative scenario: unpaired reads of length 100 with a base call error rate of 1%. We manually selected test-reference pairs of RefSeq complete Ribovirus genomes at RdRP aa identities 100%, 95% … 20%, generating simulated length-100 reads at uniformly-distributed random locations in the test genome with a mean coverage of 1000x. For `bowtie2`, the complete reference genome was used as a reference; for `DIAMOND` the reference was the translated amino acid sequence of the RdRP gene ( 400aa), which was identified by aligning to the Wolf18 dataset. These choices model the coronavirus pan-genome used as a `bowtie2` query and the rdrp1 protein reference used as a `DIAMOND` query, respectively. Sensitivity was assessed as the fraction of reads aligned to the reference. With `bowtie2`, the number of unmapped reads reflects a combination of lack of alignment sensitivity and divergence in gene content as some regions of the genome may lack homology to the reference. With `DIAMOND`, the number of unmapped reads reflects a combination of lack of alignment sensitivity and the fraction of the genome which is not RdRP, which varies by genome length 1g. They show that the fraction of aligned reads by `bowtie2` drops to around 2% to 4% at 90% RdRP aa identity, and maps no reads for most of the lower identity test-reference pairs. `DIAMOND` maps around 5% to 10% of reads down to 50% RdRP aa identity, then less than 1% at lower identities; around 30% to 35% is the lower limit of practical detection.

### 1.2 Defining viral pangenomes and the SRA search space

#### 1.2.1 Nucleotide search pangenomes

To create a collection of viral pangenomes, a comprehensive set of complete and partial genomes representing the genetic diversity of each viral family, we used two approaches.

For *Coronaviridae*, we combined all RefSeq (n = 64) and GenBank (n = 37,451) records matching the NCBI Nucleotid server query `“txid11118[Organism:exp]”` (date accessed: June 1st 2020). Sequences <200 nt were excluded as well as sequences identified to contain non-CoV contaminants during preliminary testing (such as plasmid DNA or ribosomal RNA fragments). Remaining sequences were clustered at 99% identity with `UCLUST` [62] and masked by `Dustmasker` [63] with `--window 30` and `--window 64`. The final query contained 10,101 CoV sequences (accessions in Extended Table 1a, masked coordinates in Extended Table 1b). `SeqKit` (v.0.15) was used for working with fasta files [64].

For all other vertebrate viral family pangenomes, RefSeq sequences (n = 2,849) were downloaded from the NCBI

Nucleotide server with the query `“Viruses[Organism] AND srcdb_refseq[PROP] NOT wgs[PROP] NŨT cellular organisms[ORGN] NOT AC_000001:AC_999999[PACC] AND (“vhost human”[Filter] AND “vhost vertebrates”[Filter])”` (date accessed: May 17th 2020). Retroviruses (n = 80) were excluded as preliminary testing yielded excessive numbers of alignments to transcribed endogenous retroviruses. Each sequence was annotated with its taxonomic family according to its RefSeq record; those for which no family was assigned by RefSeq (n = 81) were designated as ”unknown”.

The collection of these pangenomes was termed `cov3m`, and was the sequence reference used for this study.

#### 1.2.2 Amino acid viral RNA-dependent RNA polymerase search panproteome

For the translated-nucleotide search of viral RNA-dependent RNA polymerase (RdRP; hereinafter viral RdRP is implied) we combined sequences from several sources. 1) The ‘wolf18‘ collection is a curated snapshot (ca. 2018) of RdRP from GenBank ([21] accessed: ftp://ftp.ncbi.nlm.nih.gov/pub/wolf/_suppl/rnavir18/RNAvirome. S2.afa 2) The ‘wolf20‘ collection is RdRPs from assembled from marine metagenomes ([22] accessed: ftp://ftp.ncbi.nlm.nih.gov/pub/wolf/_suppl/yangshan/gb_rdrp.afa) 3) All viral GenBank protein sequences were aligned with `diamond --ultra-sensitive` against the combined ‘wolf18‘ and ‘wolf20‘ sequences (E-value <1e-6). These produced local alignments which contained truncated RdRP, so each RdRP-containing GenBank sequence was then re-aligned to the ‘wolf18’ and ‘wolf20‘ collection to “trim“ them to ‘wolf‘ RdRP boundaries. 4) The above algorithm was also applied to all viral GenBank nucleotide records to capture additional RdRP not annotated as such by GenBank. A region of HCV capsid protein shares similarity to HCV RdRP, sequences annotated as HCV-capsid were therefore removed. Eight novel coronavirus RdRP sequences identified in a pilot experiment were added manually. The combined RdRP sequences from the above collections were clustered (uclust) at 90% amino acid identity and the resulting representative sequences (centroids, N = 14 653) used as the rdrp1 search query.

In addition, we added Deltavirus antigen proteins from NC_001653, M21012, X60193, L22063, AF018077, AJ584848, AJ584847, AJ584844, AJ584849, MT649207, MT649208, MT649206, NC_040845, NC_040729, MN031240, MN031239, MK962760, MK962759, and eight additional homologs we identified in a pilot experiment.

#### 1.2.3 SRA search space and queries

To run `Serratus`, a target list of SRA run accessions is required. We defined eleven (not-mutually exclusive) queries as our search space which were named human, mouse, mammal, vertebrate, invertebrate, eukaryotes, prokaryotes/others, bat (including genomic sequences), virome, metagenome and mammalian genome (Extended Table 1 c). Our search was restricted to Illumina sequencing technologies and to RNA-seq, meta-genomic, and meta-transcriptome library types for these organisms (except for mammalian genome query which was genome or exome). Prior to each `Serratus` deployment, target lists were depleted of accessions already analysed. Reprocessing of a failed accession was attempted at least twice. In total, we aligned 3,837,755/4,059,695 (94.5%) of the runs in our nucleotide-pangenome search (ca. May 2020) and 5,686,715/5,780,800 (98.37%) of the runs in our translated-nucleotide RdRP search (ca. January 2021).

### 1.3 User interfaces for the Serratus databases

We implemented an on-going, multi-tiered release policy for code and data generated by this study, as follows. All code, electronic notebooks and raw data is immediately available at https://github.com/ababaian/serratus and on the `s3://serratus-public/` bucket, respectively. Upon completion of a project milestone, a structured data-release is issued containing raw data into our viral data warehouse `s3://lovelywater/`. For example, the .bam nucleotide alignment files from 3.84 million SRA runs are stored in `s3://lovelywater/bam/X.bam`; the protein .summary files are in `s3://lovelywater/psummary/X.psummary`, where `X` is a SRA run accession. These FAIR and structured releases enable downstream and third-party programmatic access to the data.

Summary files for every searched SRA dataset are parsed into a publicly accessible AWS Relational Database (RDS) instance which can be queried remotely via any `PostgreSQL` client. This enables users and programs to perform complex operations such as retrieving summaries and meta-data for all SRA runs matching a given reference sequence with above a given classifier score threshold. For example, one can query for all records containing at least 20 aligned reads to Hepatitis Delta Virus (NC_001653.2) and the associated host taxonomy for the corresponding SRA datasets:

~~~
  SELECT genbank_id, run_id, tax_id, n_reads
  FROM nsequence
  JOIN srarun ON (nsequence.run_id = srarun.run)
  WHERE n_reads >= 20
~~~

For users unfamiliar with SQL we developed `Tantalus` (https://github.com/serratus-bio/tantalus, an R programming-language package which directly interfaces the `Serratus` PostgreSQL database to retrieve summary information as data-frames. `Tantalus` also offers functions to explore and visualise the data.

Finally, the `Serratus` data can be explored via a graphical web interface by accession, virus, or viral family at https://serratus.io/explorer. Under the hood, we developed a REST API to query the database from the website. The website uses React+D3.js to serve graphical reports with an overview of viral families found in each SRA accession matching a user query.

All four data access interfaces are under ongoing development, receiving community feedback via their respective GitHub issue trackers to facilitate the translation of this data collection into an effective viral discovery resource. Documentation for data access methods is available at https://serratus.io/access.

### 1.3.1 Geo coding BioSamples

To generate the map in figure 1c, we parsed and extracted geographic information from all 16 million BioSample XML submissions. Geographic information is either in the form of coordinates (latitude/longitude) or free-form text (e.g. ”France”, ”Great Lakes”). For each BioSample, coordinate extraction was attempted using regular expressions. If that failed, text extraction was attempted using a manually curated list of keywords that capture BioSample attribute names likely to contain geographic information. If that failed, then we were unable to extract geographic information for that BioSample. Geocoding the text to coordinates was done using Amazon Location Service on a reduced set of distinct filtered text values (52,028 distinct values from 2,760,241 BioSamples with potential geographic text). BioSamples with geocoded coordinates were combined with BioSamples with submitted coordinate information to form a set of 5,325,523 geospatial BioSamples. This is then cross-referenced with our subset of SRA accessions with an RdRP match to generate the figure.

All intermediate and resulting data from this step is stored on the SQL database described in 1.3. Development work is public at https://github.com/serratus-bio/biosample-sql.

## 1.4 Viral alignment, assembly and annotation

Upon identificationn of CoV reads in a run from alignment, we assembled 52,772 runs containing at ? 10 reads which aligned to our CoV pan-genome or ≥ 2 reads with CoV-positive *k*-mers. ([17]). 11,120 of the resulting assemblies contained identifiable CoV contigs, of which only 4,179 (37.58%) contained full-length CoV RdRP (Extended Table 1). The discrepancy between alignment-positive, assembly-positive and RdRP-positive libraries arises due random sampling of viral reads and assembly fragmentation. In this respect, alignment or k-mer based methods are more sensitive than assembly in detecting for the presence of low-abundance viruses (genome coverage <1) with high identity to a reference sequence. Scoring libraries for genome-coverage and depth is a good predictor of ultimate assembly success (Extended Figure 3) thus, it can be used to efficiently prioritise computationally expensive assembly in the future.

### 1.4.1 DIAMOND optimisation and output

To optimise `DIAMOND` for small (< 10 Mb) databases such as the RdRP search database, we built a probabilistic hash set which stores 8-bit hash values for the database seeds, using SIMD instructions for fast probing. This index is loaded as a memory mapped file to be shared among processes and allows us to filter the query reads for seeds contained in the database, thus omitting the full construction of the query seed table. We also eliminated the overhead of building seed distribution histograms that is normally required to allocate memory and construct the query table in a single pass over the data using a deque-like data structure. In addition, query reads were not masked for simple repeats, as the search database is already masked. These features are available starting from `DIAMOND` v2.0.8 with the command line flags `--target-indexed --masking 0`. In a benchmark of 4 sets of 1 million read from a bat metagenome (ERR2756788), optimisation reduced `DIAMOND` runtime against RdRP-search from 197.96s (s.d=0.18s), to 21.29s (s.d=0.23s) per million reads, a speed-up of a factor of 9.3. This effectively reduced the computational cost of translated-nucleotide search for `Serratus` from $0.03, to $0.0042 per library.

`DIAMOND` output files (we label .pro) were specified with the command `-f 6 qseqid qstart qend qlen qstrand sseqid sstart send slen pident evalue cigar qseq_translated full_qseq full_qseq_mate`.

### 1.4.2 coronaSPAdes

RNA viral genome assembly faces several distinct challenges stemming from technical and biological bias in sequencing data. During library preparation, reverse transcription introduces 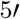 end coverage bias, and GC-content skew and secondary structures lead to unequal PCR amplification [65]. Technical bias is confounded by biological complexity such as intra-sample sequence variation due to transcript isoforms and/or to presence of multiple strains.

To address the assembly challenges specific to RNA viruses, we developed `coronaSPAdes`, described in detail in a companion manuscript [26]. In brief, `rnaviralSPAdes` and the more specialized variant, coronaSPAdes, combines algorithms and methods from several previous approaches based on `metaSPAdes` [66], `rnaSPAdes` [67] and `metaviralSPAdes` [68] with a `HMMPathExtension` step. `coronaSPAdes` constructs an assembly graph from a RNA-sequencing dataset (transcriptome, meta-transcriptome, and meta-virome are supported), removing expected sequencing artifacts such as low-complexity (poly-A / poly-T) tips, edges, single-strand chimeric loops or doublestrand hairpins [67] and subspecies-bases variation [68].

To deal with possible misassemblies and high-covered sequencing artefacts, a secondary `HMMPathExtension` step is performed to leverage orthogonal information about the expected viral genome. Protein domains are identified on all assembly graphs using a set of viral hidden Markov models (HMMs), and similar to biosyntheticSPAdes [69], `HMMPathExtension` attempts to find paths on the assembly graph which pass through significant HMM matches in order.

`coronaSPAdes` is bundled with the Pfam SARS-CoV-2 set of HMMs [70], although these may be substituted by the user. This latter feature of `coronaSPAdes` was utilized for HDV assembly, where the HMM model of HDAg, the Hepatitis Delta Antigen, was used instead of Pfam SARS-CoV-2 set. Note that despite the name, the HMMs from this set are quite general, modeling domains found in all coronavirus genera in addition to RdRP, which is found in many RNA virus families. Hits from these HMMs cover most bases in most known coronaviruse genomes, enabling the recovery of strain mixtures and splice variants.

### 1.4.3 Micro-assembly of RdRP-aligned reads

Reads aligned by `DIAMOND` in the translated-nucleotide RdRP search are stored in the `.pro` alignment file. All sets of mapped reads (3,379,127 runs) were extracted, and each non-empty set was assembled with `rnaviralSPAdes` [26] using default parameters. This process is referred to as *micro-assembly* since a collection of `DIAMOND` hits is orders of magnitude smaller than the original SRA accession (40±534 KB compressed size, ranging from a single read up to 53 MB). Then `bowtie2` [59] (default parameters) was used to align the `DIAMOND` read hits of an accession back to the micro-assembled contigs of that accession. `Palmscan --hicon` was run on micro-assembled contigs, resulting in high-confidence palmprints for 337,344 contigs. Finally `mosdepth` [71] was used to calculate a coverage pileup for each palmprint hit region within micro-assembled contigs.

### 1.4.4 Classification of assembled RdRP sequences

Our methods for RdRP classification are described and validated in a companion paper [19]. Briefly, we defined a barcode sequence, the polymerase palmprint (PP), as a ~100 aa segment of the RdRP palm sub-domain delineated by well-conserved catalytic motifs. We implemented an algorithm, `Palmscan`, to identify palmprint sequences and discriminate RdRPs from reverse transcriptases. The combined set of RdRP palmprints from public databases and our assemblies were classified by clustering into operational taxonomic units (OTUs) at 90%, 75% and 40% identity, giving species-like, genus-like and family-like clusters (sOTUs, gOTUs and fOTUs), respectively. Tentative taxonomy of novel OTUs was assigned by aligning to palmprints of named viruses and taking a consensus of the top hits above the identity threshold for each rank.

### 1.4.5 Quality control of assembled RdRP sequences

Our goal was to identity novel viral RdRP sequences and novel sOTUs in SRA libraries. From this perspective, we considered the following to be erroneous to varying degrees: sequences which are (a) not polymerases, (b) not viral, (c) with differences due to experimental artefacts, or (d) with sufficient differences to cause a spurious inference of a novel sOTU. We categorised potential sources of such errors and implemented quality control procedures to identify and mitigate them, as follows.

Point errors are single-letter substitution and indel errors which may be caused by PCR or sequencing *per se*. Random point errors are not reproduced in multiple non-PCR duplicate reads and are unlikely to assemble because such errors almost always induce identifiable structures in the assembly graph (tips and bubbles) which are pruned during graph simplification. In rare cases, a contig may contain a read with random point errors. Such contigs will have low coverage ~1, and we therefore recorded coverage as a QC metric and assessed whether low-coverage assemblies were anomalous compared to high-coverage assemblies by measures such as the frequencies with which they are reproduced in multiple libraries compared to exactly one library, finding no noticeable difference when coverage is low.

Chimeras of polymerases from different species could arise from PCR amplification or assembly. We used the UCHIME2 algorithm [72] to screen assembled palmprint sequences, finding no high-scoring putative chimeras. Mosaic sequences formed by joining a polymerase to unrelated sequence would either have an intact palmprint, in which case the mosaic would be irrelevant to our analysis, or would be rejected by `Palmscan` due to the lack of delimiting motifs.

Reverse transcriptases (RTs) are homologous to RdRP. Retroviral insertions into host genomes induce ubiquitous sequence similarity between host genomes and viral RdRP. `Palmscan` was designed to discriminate RdRP from sequences of RT origin. Testing on a large decoy set of non-RdRP sequences with recognisable sequence similarity showed that the `Palmscan` false discovery rate for RdRP identification is 0.001. We estimated the probability of false positives matches in unrelated sequence by generating sufficient random nucleotide and amino acid sequences to show that the expected number of false positive palmprint identifications is zero in a dataset of comparable size to our assemblies. We also regard the low observed frequency of palmprints in DNA WGS data (in 2.6 Pbp or 25.8% of reads, accounted for 100 known palmprints and 95 novel palmprints or 0.13% of the total identified) as a *de facto* confirmation of the low probability false positives in unrelated sequence.

Endogenous viral elements (EVEs, i.e. insertions of viral sequence into host genomes which are potentially degraded and non-functional) cannot be distinguished from viral genomes on the basis of the palmprint sequence alone. To assess the frequency of EVEs in our data, we re-assembled 890 randomly-chosen libraries yielding one or more palmprints using all reads, extracted the 23 530 resulting contigs with a positive palmprint hit by `Palmscan`, and classified them using Virsorter2 [73]. Of these contigs, 11,914 were classified as viral, confirming the `Palmscan` identification; 49 as Viridiplantae (green plants); 46 as Metazoa; 25 as Fungi and the remainder were unclassified. Thus, 120/12034 = 1% of the classified contigs were predicted as non-viral, suggesting that the frequency of EVEs in the reported palmprints is ~1%.

### 1.4.6 Annotation of CoV assemblies

Accurate annotation of CoV genomes is challenging due to ribosomal frameshifts and polyproteins which are cleaved into maturation proteins [74], and thus previously-annotated viral genomes offer a guide to accurate gene-calls and protein functional predictions. However, while many of the viral genomes we were likely to recover would be similar to previously-annotated genomes in Refseq or GenBank, we anticipated that many of the genomes would be taxonomically distant from any available reference. To address these constraints, we developed an annotation pipeline called `DARTH` [75]^1^ which leverages both reference-based and *ab initio* annotation approaches.

In brief, `DARTH` consists of the following phases: standardise the ordering and orientation of assembly contigs using conserved domain alignments, perform reference-based annotation of the contigs, annotate RNA secondary structure, *ab intio* gene-calling, generate files for aiding assembly and annotation diagnostics, and generate a master annotation file. It is important to put the contigs in the “expected” orientation and ordering to facilitate comparative analysis of synteny and as a requirement for genome deposition. To perform this standardisation, `DARTH` generates the six-frame translation of the contigs using the `transeq` [76] and uses `HMMER3` [77] to search the translations for Pfam domain models specific to CoV [70]. DARTH compares the Pfam accessions from the HMMER alignment to the NCBI SARS-CoV-2 reference genome (NCBI Nucleotide accession NC_045512.2) to determine the correct ordering and orientation, and produces an updated assembly FASTA file. `DARTH` performs reference-based annotation using `VADR` [78], which provides a set of genome models for all CoV RefSeq genomes [79]. `VADR` provides annotations of gene coordinates, polyprotein cleavage sites, and functional annotation of all proteins. `DARTH` supplements the `VADR` annotation by using `Infernal` [80] to scan the contigs against the SARS-CoV-2 Rfam release [81] which provides updated models of CoV 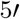 and 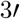 untranslated regions (UTRs) along with stem-loop structures associated with programmed ribosomal frame-shifts. While `VADR` provides reference-based gene-calling, `DARTH` also provides *ab initio* gene-calling by using `FragGeneScan` [82], a frameshift-aware gene caller. `DARTH` also generates auxiliary files which are useful for assembly quality and annotation diagnostics, such as indexed BAM files created with `SAMtools` [83] representing self-alignment of the trimmed reads to the canonicalized assembly using `bowtie2` [59], and variant-calls using `bcftools` from `SAMtools`. `DARTH` generates these files so that the can be easily loaded into a genome browser such as `JBrowse` [84] or IGV [85]. As the final step `DARTH` generates a single Generic Feature Format (GFF) 3.0 file [86] containing combined set of annotation information described above, ready for use in a genome browser, or for submitting the annotation and sequence to a genome repository.

### 1.4.7 Phage assembly

Each metagenomic dataset was individually *de novo*-assembled using `MEGAHIT` [87], and filtered to remove contigs smaller than 1 kbp in size. ORFs were then predicted on all contigs using `Prodigal` v2.6.3 [88] with the following parameters: `-m -p meta`. Predicted ORFs were initially annotated using `USEARCH` [62] to search all predicted ORFs against `UniProt` [89], `UniRef90` and `KEGG` [90]. Sequencing coverage of each contig was calculated by mapping raw reads back to assemblies using `Bowtie2` [59]. Terminase sequences from Al-Shayeb *et al.* [45] were clustered at 90% amino acid identity to reduce redundancy using `CD-HIT` [91], and HMM models were built with `hmmbuild` (from the `HMMER3` suite [77]) from the resulting set. Terminases in the assemblies from `Serratus` were identified using `hmmsearch`, retaining representatives from contigs greater than 140 kbp in size. Some examples of prophage and large phages that did not co-cluster with the sequences from Al-Shayeb *et al.*, were also recovered because they were also present in a sample that contained the expected large phages. The terminases were aligned using `MAFFT` v.7.407 [92] and filtered by TrimAL [93] to remove columns comprised of more than 50% gaps, or 90% gaps, or using the automatic `gappyout` setting to retain the most conserved residues. Maximum likelihood trees were built from the resulting alignments using `IQTREE` v.1.6.6 [94].

### 1.4.8 Deploying the assembly and annotation workflow

The `Serratus` search for known or closely related viruses identified 37,131 libraries (14,304 by nucleotide and 23,898 by amino acid) as potentially positive for CoV (score ≥20 and ≥10 reads). To supplement this search we also employed a recently developed index of the SRA called STAT [17] with which identified an additional 18,584 SRA datasets not in the defined SRA search space. The STAT BigQuery was `WHERE tax_id=11118 AND total_count` >1 accessed on June 24th 2020.

We used AWS Batch to launch thousands of assemblies of NCBI accessions simultaneously. The workflow consists of four standard parts: a job queue, a job definition, a compute environment, and finally, the jobs themselves. A CloudFormation template^2^ was created for building all parts of the cloud infrastructure from the command line. The job definition specifies a Docker image, and asks for 8 virtual CPUs (vCPUs, corresponding to threads) and 60 GB of memory per job, corresponding to a reasonable allocation for `coronaSPAdes`. The compute environment is the most involved component. We set it to run jobs on cost-effective Spot instances (`optimal` setting) with an additional cost-optimization strategy (`SPOT_CAPACITY_OPTIMIZED` setting), and allowing up to 40,000 vCPUs total. In addition, the compute environment specifies a launch template which, on each instance, i) automatically mounts an exclusive 1 TB EBS volume, allowing sufficient disk space for several concurrent assemblies, and ii) downloads the 5.4 GB CheckV [95] database, to avoid bloating the Docker image.

The peak AWS usage of our Batch infrastructure was ~28,000 vCPUs, performing ~3,500 assemblies simultaneously. A total of 46,861 accessions out of 55,715 were assembled in a single day. They were then analysed by two methods to detect putative CoV contigs. The first method is CheckV [95], followed selecting contigs associated to known CoV genomes. The second method is a custom script^3^ that parses `coronaSPAdes` BGC candidates and keeps contigs containing CoV domain(s). For each accession, we kept the set of contigs obtained by the first method (CheckV) if it is non-empty, and otherwise we kept the set of contigs from the second method (BGC).

A majority (76%) of the assemblies were discarded for one of the following reasons: i) no CoV contigs were found by either filtering method, ii) reads were too short to be assembled, iii) Batch job or SRA download failed, or iv) `coronaSPAdes` ran out of memory. A total of 11,120 assemblies were considered for further analysis.

The average cost of assembly was between $0.30-$0.40 per library, varying depending on library-type (RNA-seq versus metagenomic). This places an estimate of 46-95 fold higher cost for assembly alone compared to a cost of $0.0042 or $0.0065 for an alignment based search.

## 1.5 Taxonomic and phylogenetic analyses

### 1.5.1 Taxonomy prediction for coronavirus genomes

We developed a module, SerraTax, to predict taxonomy for CoV genomes and assemblies (https://github.com/ababaian/serratus/tree/master/containers/serratax). `SerraTax` was designed with the following requirements in mind: provide taxonomy predictions for fragmented and partial assemblies in addition to complete genomes; report best-estimate predictions balancing over-classification and under-classification (too many and too few ranks, respectively); and assign an NCBI Taxonomy Database [96] identifier (`TaxID`).

Assigning a best-fit `TaxID` was not supported by any previously published taxonomy prediction software to the best of our knowledge; this requires assignment to intermediate ranks such as sub-genus and ranks below species (commonly called strains, but these ranks are not named in the Taxonomy database), and to unclassified taxa, e.g. `TaxID 2724161, unclassified Buldecovirus`, in cases where the genome is predicted to fall inside a named clade but outside all named taxa within that clade.

`SerraTax` uses a reference database containing domain sequences with `TaxIDs`. This database was constructed as follows. Records annotated as CoV were downloaded from `UniProt` [89], and chain sequences were extracted. Each chain name, e.g. `Helicase`, was considered to be a separate domain. Chains were aligned to all complete coronavirus genomes in `GenBank` using `UBLAST` [62] to expand the repertoire of domain sequences. The reference sequences were clustered using `UCLUST` [62] at 97% sequence identity to reduce redundancy.

For a given query genome, open reading frames (ORFs) are extracted using the `EMBOSS getorf` software [76]. ORFs are aligned to the domain references and the top 16 reference sequences for each domain are combined with the best-matching query ORF. For each domain, a multiple alignment of the top 16 matches plus query ORF is constructed on the fly by `MUSCLE` [97] and a neighbour-joining tree is inferred from the alignment, also using `MUSCLE`. Finally, a consensus prediction is derived from the placement of the ORF in the domain trees. Thus, the presence of a single domain in the assembly suffices to enable a prediction; if more domains are present they are combined into a consensus.

### 1.5.2 Taxonomic assignment by phylogenetic placement

To generate an alternate taxonomic annotation of an assembled genome, we created a pipeline based on phylogenetic placement, `SerraPlace`.

To perform phylogenetic placement, a reference phylogenetic tree is required. To this end, we collected 823 reference amino acid RdRP sequences, spanning all *Coronaviridae*. To this set we added an outgroup RdRP sequence from the Torovirus family (NC_007447). We clustered the sequences to 99% identity using USEARCH ([62], `UCLUST` algorithm, `v11.0.667`), resulting in 546 centroid sequences. Subsequently we performed multiple sequence alignment on the clustered sequences using `MUSCLE` ([97], v3.8.31). We then performed maximum likelihood tree inference using `RAxML-NG` ([98], `PROTGTR+FO+G4, v0.9.0`), resulting in our reference tree.

To apply SerraPlace to a given genome, we first use HMMER ([77], v3.3) to generate a reference HMM, based on the reference alignment. We then split each contig into ORFs using `esl-translate`, and use `hmmsearch` (p-value cutoff 0.01) to identify those query ORFs that align with sufficient quality to the previously generated reference HMM. All ORFs that pass this test are considered valid input sequences for phylogenetic placement. Subsequently, we use `EPA-ng` ([99], `v0.3.7`) to place each sequence on the RdRP reference tree. This produces a set of likely placement locations on the tree, with an associated likelihood weight. We then use `Gappa` ([100], v0.6.1) to assign taxonomic information to each query, using the taxonomic information for the reference sequences. `Gappa` assigns taxonomy by first labelling the interior nodes of the reference tree by a consensus of the taxonomic labels of all descendant leaves of that node. If 66% of leaves share the same taxonomic label up to some level, then the internal node is assigned that label. Then, the likelihood weight associated with each sequence is assigned to the labels of internal nodes of the reference tree, according to where the query was placed.

From this result, we select that taxonomic label that accumulated the highest total likelihood weight as the taxonomic label of a sequence. Note that multiple ORFs of the same genome may result in a taxonomic label, in which case, we select the longest sequence as the source of the taxonomic assignment of the genome.

### 1.5.3 Phylogenetic inference

We performed phylogenetic inferences using a custom snakemake pipeline (available at https://github.com/lczech/nidhoggr), using ParGenes [101], `v1.1.2`. `ParGenes` is a treesearch orchestrator, built on top of `ModelTest-NG` [102] and `RAxML-NG`, enabling higher levels of parallelisation for a given tree search.

To infer the maximum likelihood phylogenetic trees, we performed a tree search comprising 100 distinct starting trees (50 random, 50 parsimony), as well as 1000 bootstrap searches. We used `ModelTestNG` to automatically select the best evolutionary model for any given treesearch. The pipeline also automatically produces versions of the best maximum likelihood tree annotated with Felsenstein’s Bootstrap ([103]) support values, and Transfer Bootstrap Expectation ([104]) values.

## Data availability

Archival copies of all code generated for this study is available at https://github.com/serratus-bio. Electronic notebooks for experiments are available at https://github.com/ababaian/serratus. Access to all data generated in this study can be accessed at https://serratus.io/access. Assembled genomes contigs for this study are available at https://serratus.io/access pending deposition into public repositories.

## Acknowledgments

The `Serratus` project is an initiative of the hackseqRNA genomics hackathon (https://www.hackseq.com). We would like to thank the many contributors for code snippets and bioinformatic discussion; E. Erhan, J. Chu, S. Jackman, I. Birol, K. Wellman, O. Fornes, C. Xu, M. Huss, K. Ha, M. Krzywinski, E. Nawrocki, R. McLaughlin, C. Morgan-Lang, C. Blumberg, and the J. Brister lab. A. Rodrigues, S. McMillan, V. Wu, C. Kennett, K. Chao, and N. Pereyaslavsky for AWS support. We would also like to thank the J. Joy lab, G. Mordecai, J. Taylor, S. Roux, N. Kyrpides, T. Reddy, L. Bergner, R. Orton, and D. Streicker for virology discussions. We are grateful to the entire team managing the NCBI SRA. TA is grateful for the Advanced Research Computing resource at the University of British Columbia. PB was financially supported by the Klaus Tschira Foundation, RC by ANR Transipedia, Inception and PRAIRIE grants (PIA/ANR-16-CONV-0005, ANR-18-CE45-0020, ANR-19-P3IA-0001), MdlP by Ministerio de Economía y Competitividad of Spain and FEDER grant (BFU2017-87370-P), AK and DM were supported by the Russian Science Foundation (grant 19-14-00172) and computation was carried out in part by Resource Centre “Computer Centre of SPbU”. AK and DM are grateful to Saint Petersburg State University for the overall support of this work. Project support and computing resources were kindly provided by the University of British Columbia Community Health and Wellbeing Cloud Innovation Centre, powered by AWS.

Special thanks to our patient and understanding partners.

## Author contributions

AB conceived and led the study. AB and JT designed and implemented the `Serratus` architecture. AB and RCE constructed the virus pangenomes and RdRP query. RCE developed the `SerraTax` and `Summarizer` modules. PB developed the `SerraPlace` tree placement and taxonomy prediction code and calculated maximum likelihood trees. TA developed the `DARTH` annotation pipeline and submitted the annotated genomes to ENA. DM and AK developed the `coronaSPAdes` assembler. RC implemented the assembly pipeline, and deployed the assembly and annotation pipeline. BB optimised `DIAMOND` algorithm for RdRP search. AB, VL, and DL designed and developed https://serratus.io and the SQL server. AB and GN developed the `Tantalus R` package. AB, RCE, TA, PB, DM, MdlP, AK, and RC analysed the coronavirus, RdRP and deltavirus data. BAS and JB designed the phage panproteome, assembled phage genomes, and conducted phage phylogenetic analyses. All authors contributed to data interpretation and writing the manuscript.

## Correspondence

Correspondence should be addressed to AB.

## Reporting Summary

Does not apply.

## Extended Tables

Extended Table 1: **SRA run queries and CoV assembly table** Queries and accessions from this study. **a** SRA queries to retrieve collections of datasets. **b** Run accessions, assembly statistics and select meta-data for the 11,120 runs for which *Coronaviridae*, or *Coronaviridae*-like sequences were assembled. **c** Assignment of assembled runs to operational taxonomic units (OTUs) based on 97% identity of the RNA dependent RNA polymerase (RdRp) domain. **d** Assignment of GenBank records to RdRp OTUs. **e** Assignment of expected viral host for GenBank records based on Sequence Read Archive and JGI GOLD metadata [2, 31]. **f** Taxonomic source for RdRp containing assemblies.}

## 2 Extended Figures

**Extended Figure 1:**
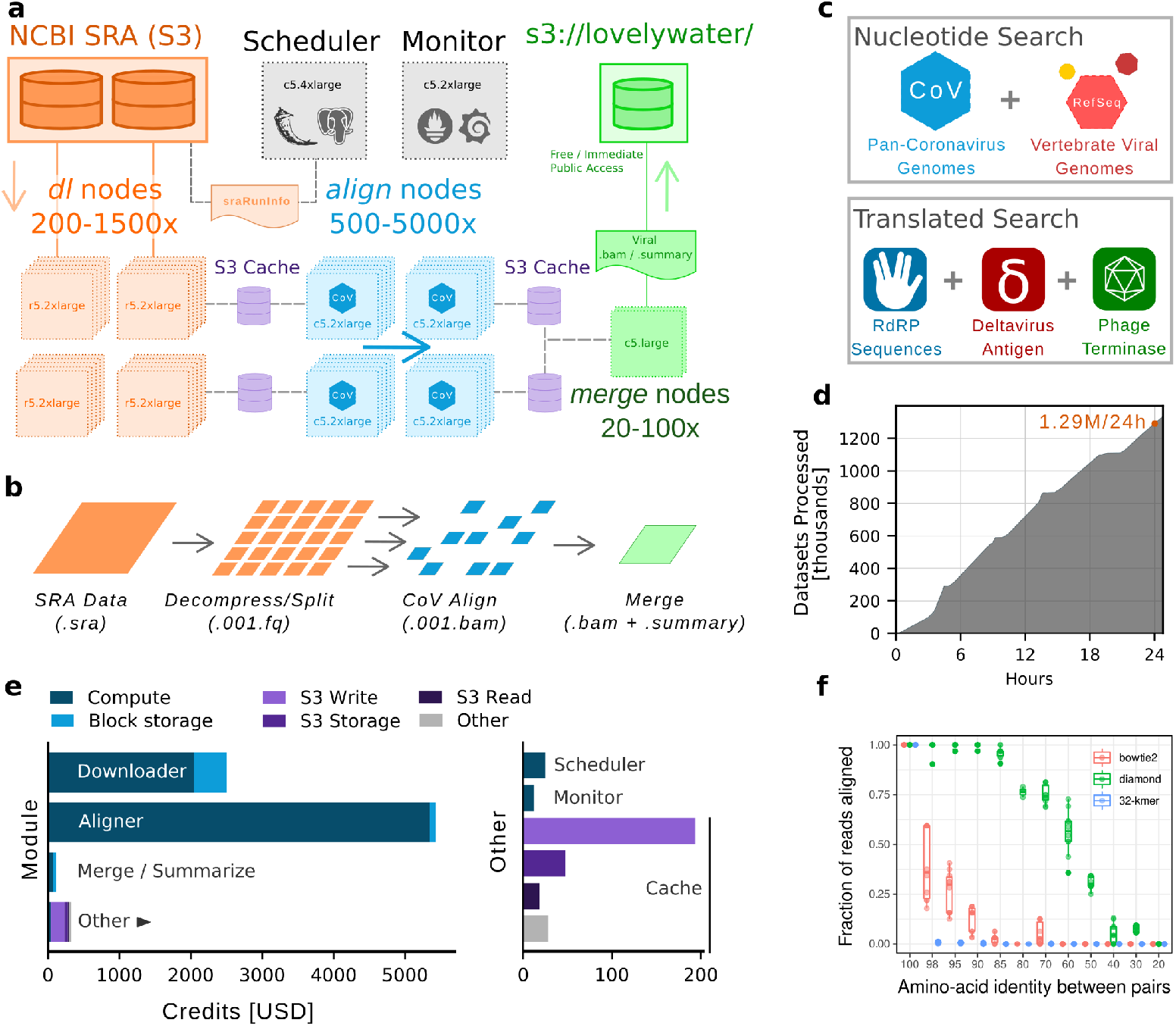
Overview of the Serratus infrastructure. **a** Schematic and data workflow (**b**) as described in the methods for sequence alignment. **c** The `align` module accepts either a nucleotide or protein sequence query. **d** A nucleotide alignment completion rate for *Serratus* shows stable and linear performance to complete 1.29 million SRA accessions in a 24-hour period and the **e** cost breakdown for this run. Compute costs between modules are an approximate comparison of CPU requirements of each step. The total average cost per completed SRA accession was $0.0062 US dollars for nucleotide search or $0.0042 for translated-nucleotide search. **f** Biological cross-validation to measure alignment sensitivity for `bowtie2` (nucleotide search), `diamond` (translated nucleotide search) or `32 kmer` for exact search. Briefly, two RdRP sequences sharing the nominal amino acid identities form a “pair”. 100 bp reads were simulated from the coding sequence of one pair and mapped onto the second pair, with the fraction of reads mapped reported. A value of 50% indicates that half the simulated reads at the given RdRP percent identity are mappable and thus detectable (see Methods).

**Extended Figure 2:**
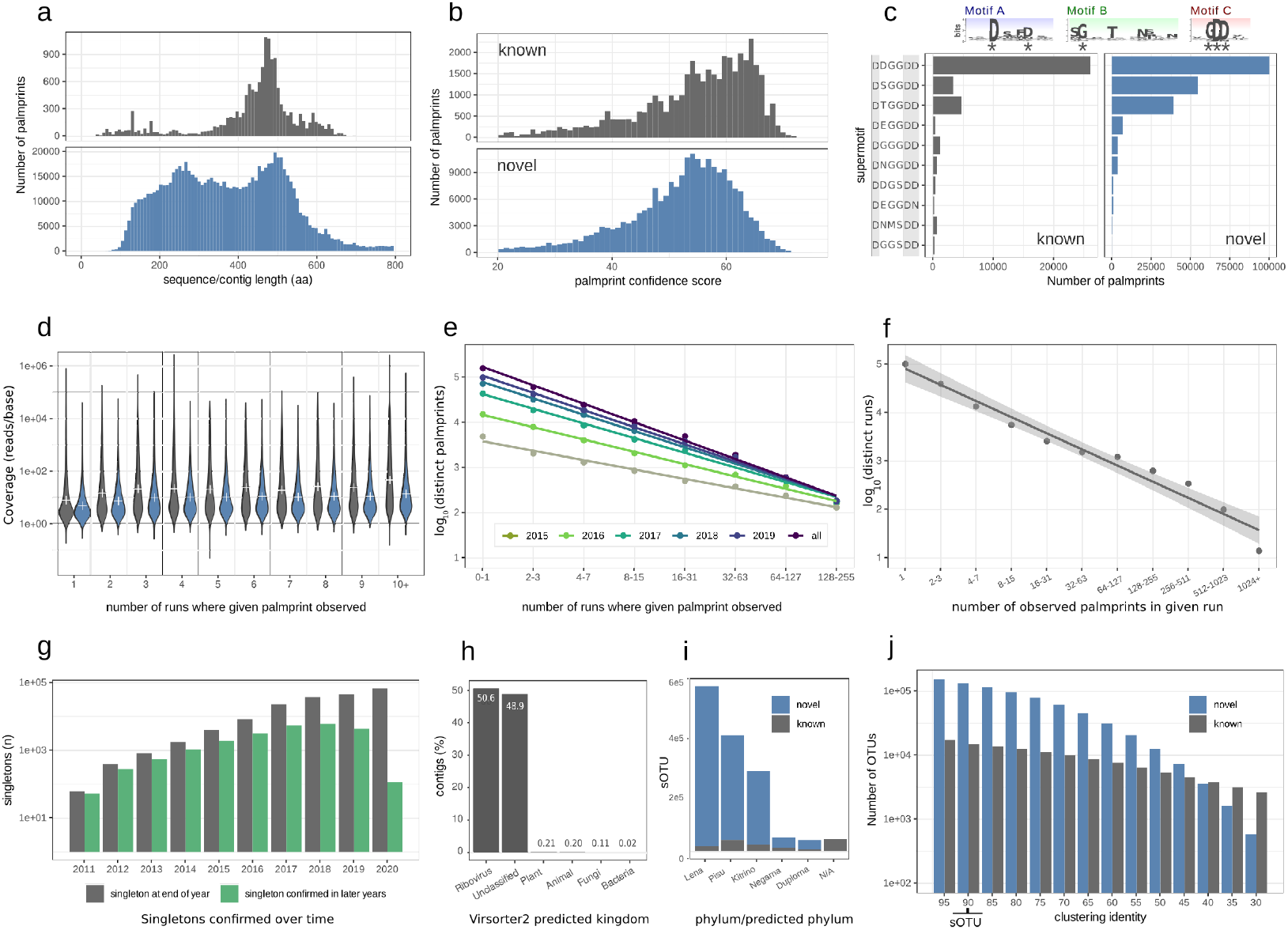
Analysis of palmprint contigs recovered by Serratus. **a** Length distribution of amino acid sequences in the rdrp1 query (upper histogram) and micro-assembled contigs (lower histogram, length=nucleotides/3). **b** Distribution of `Palmscan` confidence scores. **c** Observations of the 10 most frequent “super-motifs” (six well-conserved residues marked with asterisk) reported by `Palmscan`. **d** Distribution of coverage vs. abundance (number of runs where a given palmprint is observed), showing that palmprints have similar underlying coverage distributions at all abundances. **e** Preston plot of distinct palmprints vs. abundance exhibiting similar, approximately log-log-linear relationships to totals at end-of-year 2015 to 2019 and final totals at approx. end of 2020 (all). **f** Preston plot of number of distinct palmprints observed in a given run vs. number of runs. **g** Numbers of singletons and second observations (confirmations) at the end of each year showing that the growth in singletons is matched by a comparable growth in confirmations. **h** Kingdom predicted by `Virsorter2for` RdRP+ contigs (by `Palmscan`) obtained by full assembly of 880 randomly-chosen RdRP+ runs. **i** Number of palmprints in each phylum assigned by taxonomy (known) or predicted (novel). **j** Number of OTUs as a function of clustering identity.

**Extended Figure 3:**
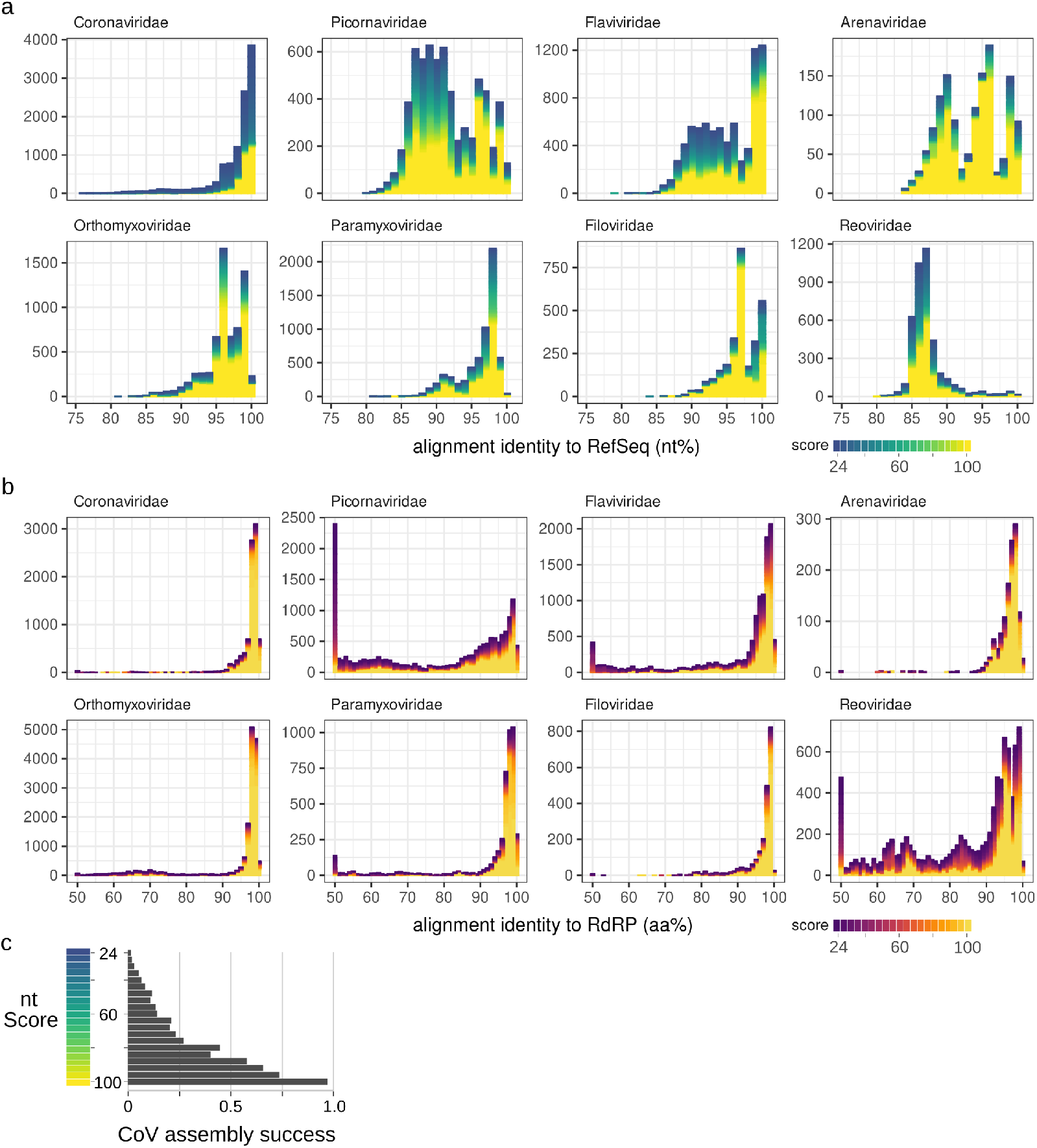
Distribution of select RNA virus families. Histogram of datasets matching select RNA viral family by (**a**) nucleotide search against RefSeq pangenomes or (**b**) translated-nucleotide search against RdRP query, binned by the average nucleotide or amino acid identity, respectively. Score (gradient colouring) function approximates pangenome/gene coverage (see methods) used for manual inspection and to prioritise assembly. Coarsely, nucleotide identity between 80-100% approximate amino acid identity of 90-100%. Interactive and queryable versions of these plots for extended virus families are available at https://serratus.io/explorer. **c** Relationship between the pangenome score function and the subsequent assembly success (defined by the presence of an RdRP+ contig) measured from 52,772 libraries with reads aligning to Coronaviridae.

**Extended Figure 4:**
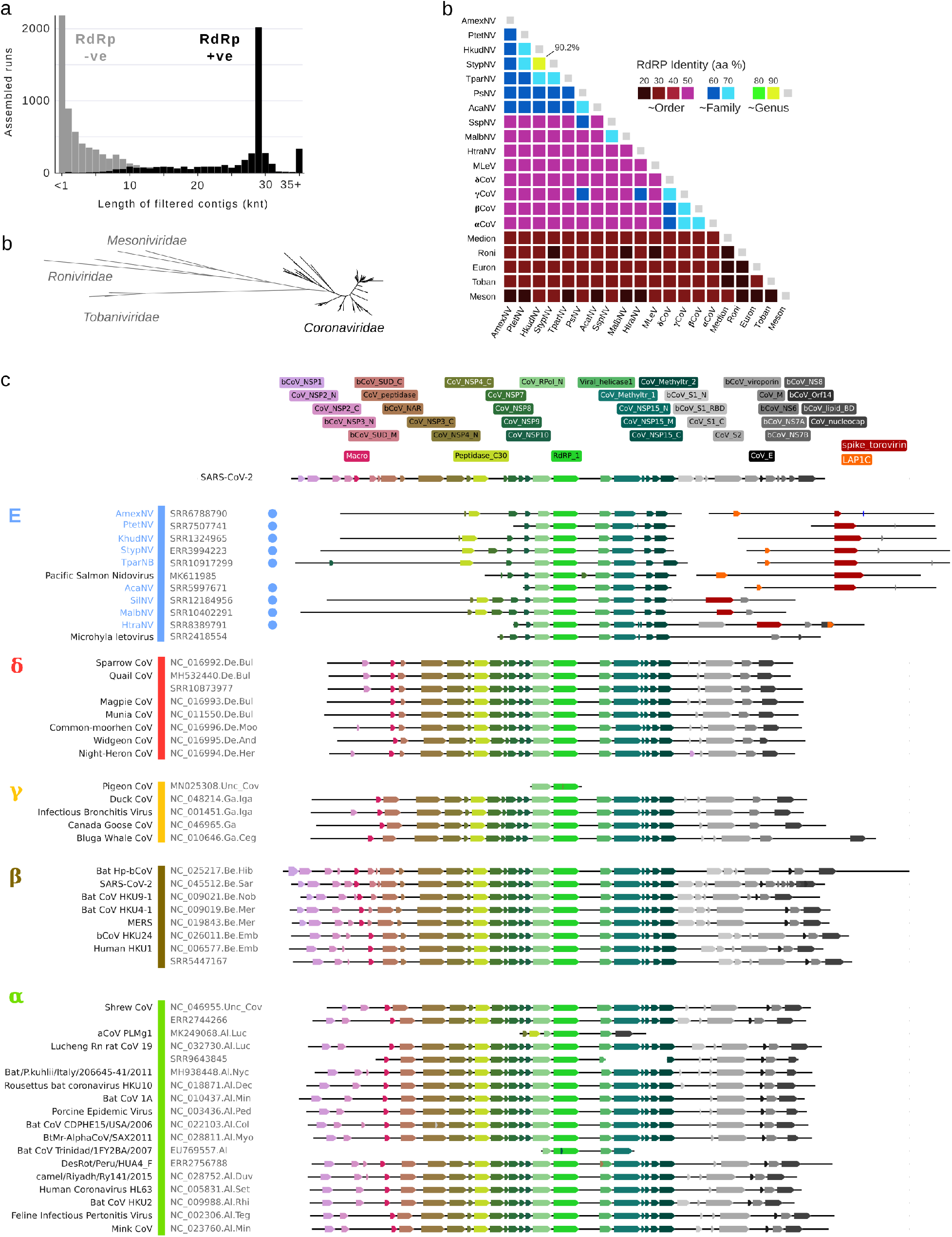
Genome organisation of *Coronaviridae* and neighbours. **a** Length distribution for 11,120 assembled contigs classified as CoV-positive, showing a peak around the typical CoV genome length, 4,179 (37.58%) of contigs also contained a match for RdRP. **b** Phylogram shown in Figure 3 showing the *Mesoniviridae*, *Tobaniviridae*, and *Roniviridae* outgroups. **c** Triangular matrix showing median RdRp sequence identities between selected *Nidovirales* and group *E* sequences. **d** Hidden Markov Model (HMM) protein domain matches from the RdRp in exemplar sequences (contigs or GenBank sequences), grouped by genus. Novel sOTUs identified in this analysis indicated by a coloured circle.

**Extended Figure 5:**
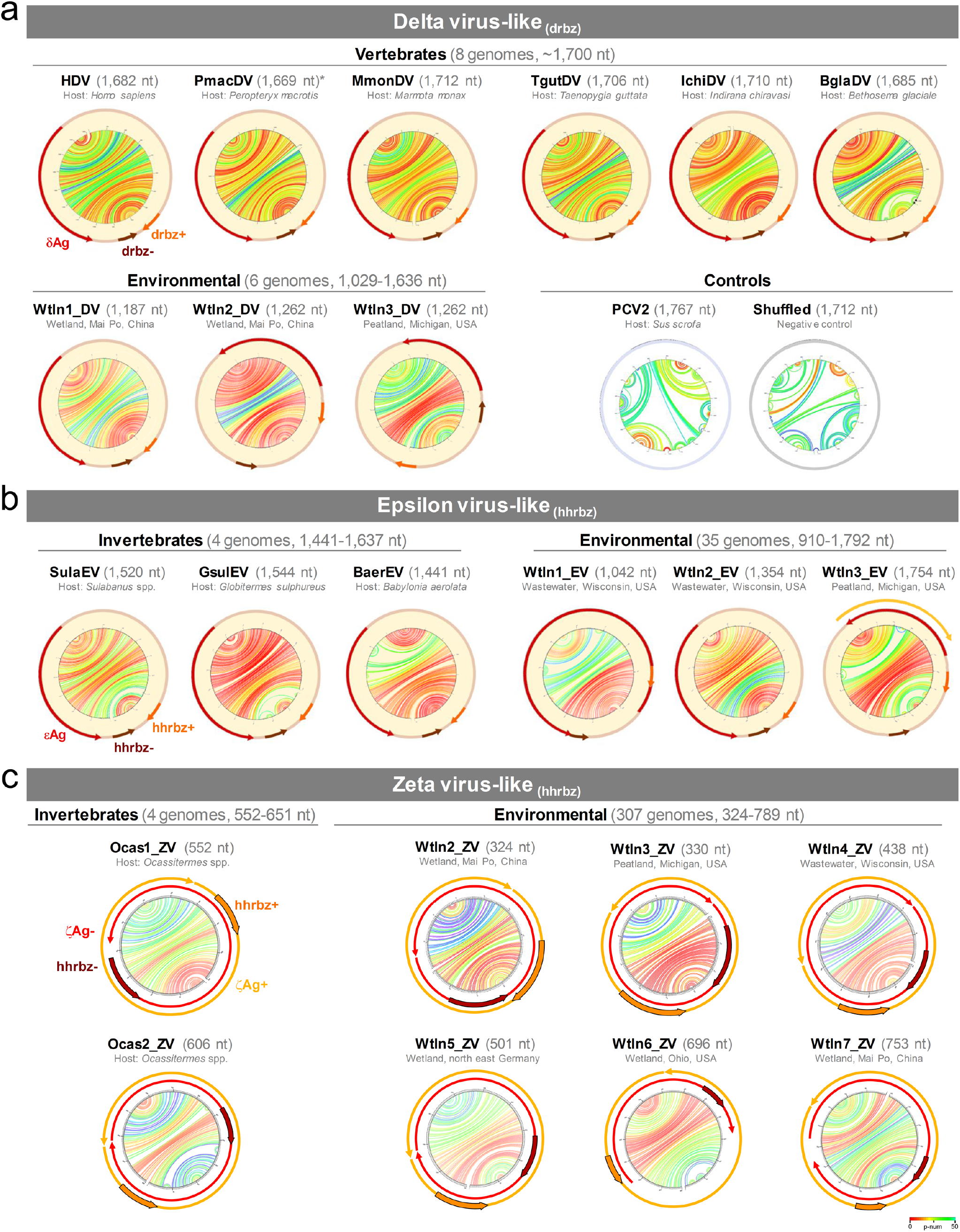
Newly characterised Deltavirus and Deltavirus-like genomes. Structure and organisation of selected examples from the 14 delta virus-, 39 epsilon virus- and 311 zeta virus-like genomes identified in our study. **a** Similar to human deltavirus (HDV), delta virus-like genomes from vertebrates (PmacDV SRR7910143; MmonDV SRR2136906; TgutDV SRR5001850; IchiDV SRR8954566 and BglaDV SRR8242383) and environmental datasets (SRR7286070 and SRR6943136) share similar predicted stable rod-like folding, a predicted ORF coding for the delta antigen (*δAg*) and a delta ribozyme (dvrbz) on each polarity. Folding of the circular DNA virus Porcine Circovirus 2 (PCV2) and a shuffled MmonDV sequence are shown as negative controls. **b** Epsilon virus-like genomes detected in invertebrates (SulaEV SRR8739608; GsulEV SRR7170939 and BaerEV SRR12300397) and environmental datasets (SRR8840728 and SRR6943136) show similar structure and organization to deltaviruses, with one or two predicted ORFs (epsilon antigen or *ϵ*Ag) and two hammerhead ribozymes (hhrbz) in equivalent genomic regions. **c** Zeta virus-like genomes detected in invertebrate (Ocassitermes sp. ZVs SRR8924823)and environmental datasets (SRR7286070, SRR6943136, SRR8840728, SRR6201737, SRR5864109 and SRR12063536) are smaller than delta and epsilon agents. Up to 90% of the zeta genomes have sizes multiple of 3 and predicted ORFs without stop codons, capable to encode endless tandem-repeated zeta antigens in both polarities (*ζ*Ag+ and *ζ*Ag− shown as yellow and red arrows, respectively). Both genomic zeta polarities keep hhrbzs (shown as arrows overlapping the ORFs) similar to the epsilon ribozymes (Extended Fig 6). Larger zeta virus-like genomes (>651 nt) were less abundant (7% of all zeta genomes) and frequently show stop codons, or their sizes are not multiple of 3.

**Extended Figure 6:**
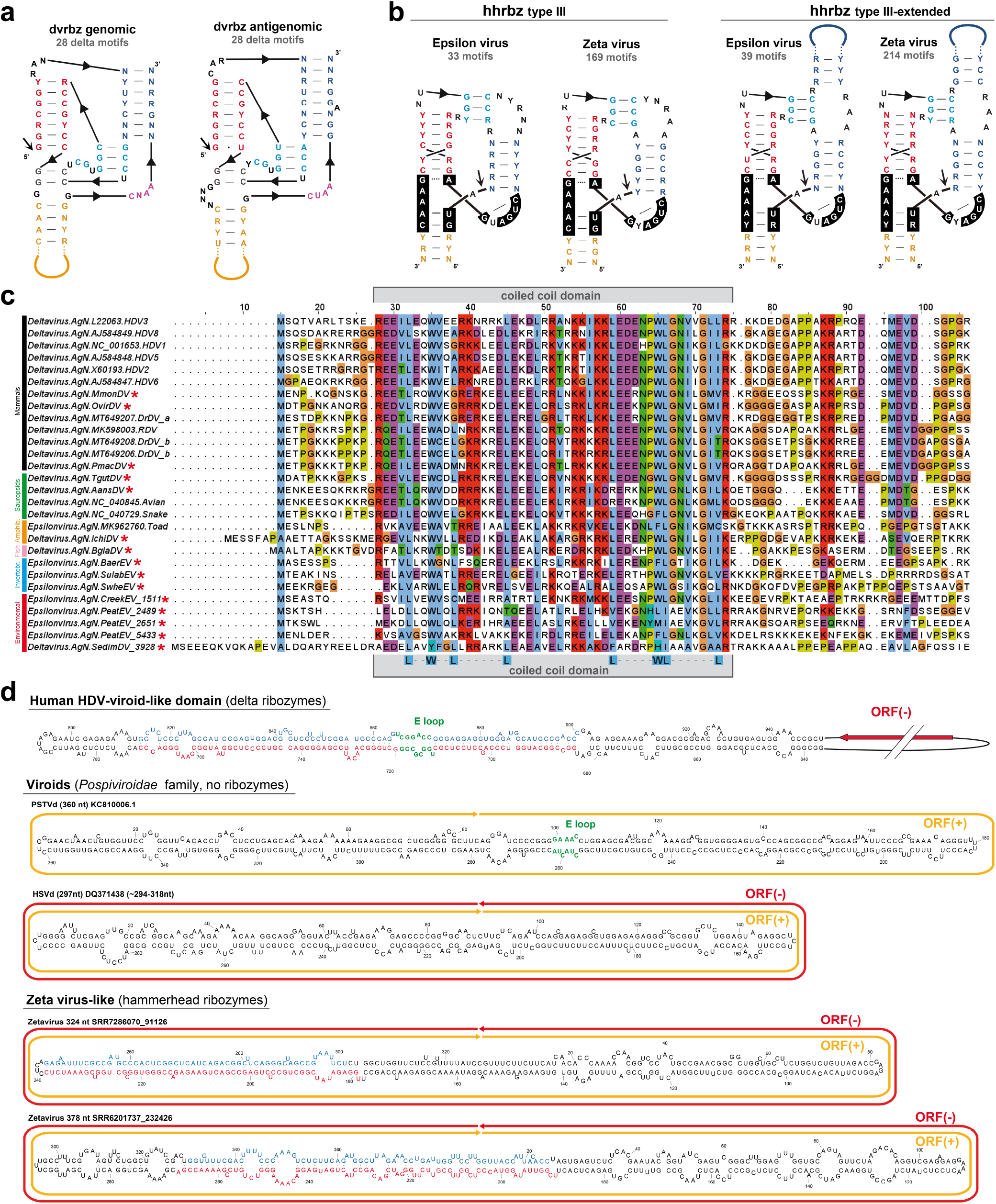
Evolutionary history of deltavirus-like agents. **a** Consensus structures (weighted nucleotide conservation threshold of 90%) of deltavirus ribozymes, including the 14 genomes described in this work.**b** Consensus structures of the two hammerhead ribozyme families (type III and extended-type III [43]) detected in epsilon and zeta agents. Most positions of epsilon and zeta motifs are sequence conserved for each ribozyme family.**c** MSA of the predicted antigen (N-term domain) from delta and epsilon agents (genomes detected in this study are indicated with a red asterisk). The antiparalel coiled-coil of the HDV is delimited with a grey box, and conserved residues involved in hydrophobic interactions are shown at the bottom [46], supporting a highly divergent connection between delta and epsilon genomes. **d** Human HDV deltavirus is known to contain a viroid-like domain related to the *Pospiviroidae* family of plant viroids. Both families of agents conserve a tertiary structure reminiscent of the E-loop 5S rRNA (nucleotides in green) and are replicated by the RNA Pol II of the host [47]. *Pospiviroids*, despite of lacking hhrbzs, share with zeta genomes a small rod structure, and in some cases, the presence of predicted endless tandem-repeat ORFs, most notably in both polarities of numerous variants of the Hop Stunt Viroid (HSVd). Whereas viroids have been historically regarded as non protein coding RNAs, our reported observations warrant further investigation.

**Extended Figure 7:**
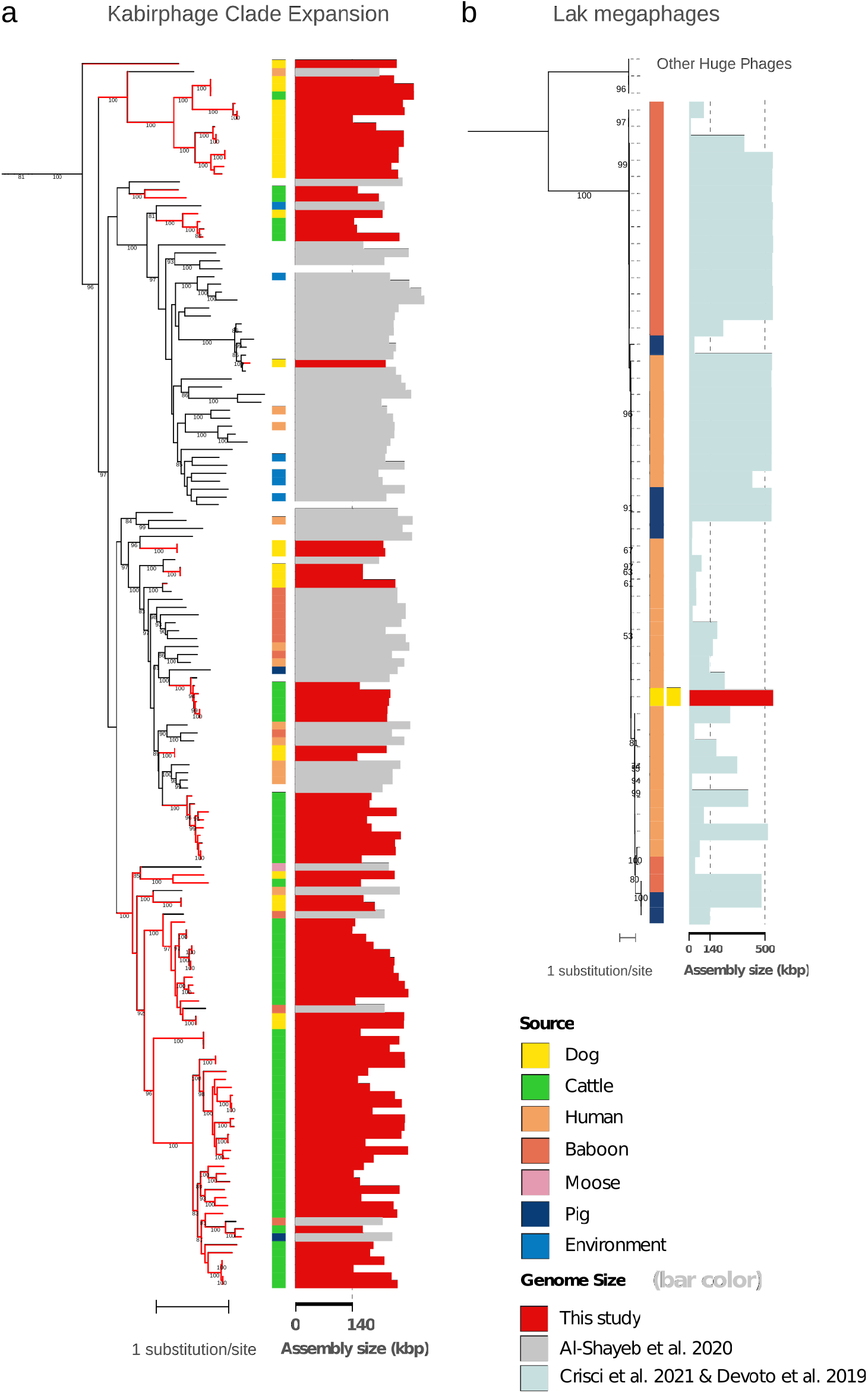
Huge phage and Lak phage detail. Expanded view of maximum likelihood terminase large subunit protein phylogenetic trees for (**a**) the expansion of the Kabirphage clade by newly recovered sequences from different animal types (colored dots). Red branches are public data recovered by Serratus, black branches indicate the previously reported genomes from [45]. **b** Publicly available Lak phage genomes [105] with sequences of two newly reconstructed complete Lak megaphage genomes. These are the first reported Lak megaphages from dogs (assembled from fecal sample metagenome reads from Allaway et al. 2020). The genomes have identical terminase sequences (at the nucleotide level) although the dogs were in different housing areas and were sampled at different times (D Allaway, personal communication).

## Footnotes

1 https://bitbucket.org/tomeraltman/darth/

2 https://gitlab.pasteur.fr/rchikhi_pasteur/serratus-batch-assembly/-/blob/master/template/template.yaml

3 https://gitlab.pasteur.fr/rchikhi_pasteur/serratus-batch-assembly/-/blob/master/stats/bgc_parse_and_extract.py

